# PFOA induced metabolic and immune perturbations in a SARS-2 infection model

**DOI:** 10.64898/2026.02.21.707085

**Authors:** Deanna N. Lanier, Dawne Rowe Haas, Mario Uchimiya, Cheryl Jones, Scott Johnson, Kaori Sakamoto, Jessie R. Chappel, Allison N. Fry, Franklin E. Leach, Jamie DeWitt, Tracey Woodlief, David A. Gaul, Erin S. Baker, Facundo M. Fernández, Stephen M. Tompkins, Arthur S. Edison

## Abstract

This study evaluates the impact of PFOA exposure on the metabolome and immune response to SARS-2 using a ferret model. Ferrets were separated into control or PFOA-exposed groups (10/mg/kg/day) and challenged with SARS-2. Longitudinal analyses encompassing clinical assessments, serological profiling, histopathology, and untargeted nuclear magnetic resonance (NMR) metabolomics revealed significant metabolic and immunological perturbations. We found prominent effects of PFOA exposure on metabolism, which resulted in altered metabolic responses to SARS-2 infection. PFOA exposure was also associated with impaired immune function, as evidenced by decreased serum IgG levels, increased viral loads, and prolonged peak infectivity. These findings underscore the consequences of PFOA exposure on host metabolism and immunity during infectious diseases.

## INTRODUCTION

Human beings are continuously exposed to a variety of chemicals through food, drinking water, and consumer products ^[1]^. Among these, per- and polyfluoroalkyl substances (PFAS) represent a globally significant group of synthetic compounds that have garnered attention due to their environmental persistence and potential adverse health effects ^[2–4]^. Perfluorooctanoic acid (PFOA), a key legacy PFAS of concern, is a particular focus of research due to its toxic properties, classification as a Group 1 carcinogen ^[5]^, extensive environmental distribution ^[4]^, and bioaccumulation in human and animal tissues ^[6]^. The chemical stability of PFOA is driven by strong carbon-fluorine bonds, which contribute to its resistance to degradation and make it a persistent contaminant in ecological and biological systems ^[6–8]^. PFAS, once believed to be chemically inert, have been demonstrated to be biologically active, interfering with normal metabolic processes and impacting key metabolic pathways ^[9, 10]^. For example, PFAS exposure has been linked to alterations in lipid ^[9, 11, 12]^, amino acid ^[11, 12]^, and carbohydrate metabolism ^[11]^, and impaired thyroid homeostasis ^[7]^, which may contribute to broader systemic health effects^[9, 10]^. Beyond metabolic and endocrine disruptions, PFOA has demonstrated significant immunotoxic potential ^[13]^. PFOA exposure has been associated with the suppression of antibody responses ^[14, 15]^, increased susceptibility to infectious diseases ^[15]^, and a greater incidence of autoimmune disorders ^[16]^. These findings suggest that PFOA can influence multiple aspects of immune regulation, though the precise mechanisms of action are not fully understood.

The global COVID-19 pandemic, caused by severe acute respiratory syndrome coronavirus 2 (SARS-2), has affected millions worldwide ^[17]^. The variability in COVID-19 outcomes, ranging from asymptomatic presentations to severe respiratory failure and death, underscores the need to investigate potential exogenous contributors to the heterogeneity in immune responses ^[18–22]^. In this context, environmental toxicants such as PFOA have emerged as critical factors to assess for their impact on health outcomes ^[23, 24]^. More specifically, recent studies suggest a link between PFAS exposure—including PFOA—and COVID-19 susceptibility ^[25–27]^ and adverse outcomes. These include higher ICU admissions ^[28]^ and increased risk of severe or fatal COVID-19 ^[27, 29, 30]^. These findings collectively suggest that PFOA exposure may worsen clinical outcomes following SARS-2 infection, though the underlying metabolic changes remain poorly defined and require investigation in controlled models. Shifts in metabolic signatures can disrupt physiological homeostasis in ways that contribute to disease development and progression, while also serving as early biomarkers that precede and predict overt pathology and/or disease states.

To address this gap, we designed a longitudinal study using ferrets as a model system. Ferrets are highly susceptible to SARS-2 and share key anatomical and physiological similarities with the human respiratory tract, making them an ideal model for studying COVID-19 pathogenesis ^[31–34]^. By utilizing untargeted metabolomics based on nuclear magnetic resonance (NMR) spectroscopy, we conducted a comprehensive analysis of metabolic changes associated with PFOA exposure and SARS-2 infection in ferret serum and tissues. This NMR-based approach provides a reproducible and non-destructive means of evaluating metabolic perturbations and enables the identification of unexpected metabolic changes ^[35–37]^. The findings from this study provide critical insights into the interaction between PFOA and COVID-19 progression.

## RESULTS

### Results Section 1: PFOA Administration Establishes Sustained Exposure Without Impacting Body Weight in Ferrets

To investigate PFOA detectability and potential toxicity in ferrets, PFOA was mixed with feed at doses of 0 mg/kg/day (control), 2 mg/kg/day (low), or 10 mg/kg/day (high) over 21 days (Figure 1A). Serum samples were collected at baseline (day -5) and at 7, 14, and 21 days post-exposure (DPE) to quantify PFOA levels using liquid chromatography-mass spectrometry (LC-MS). Peak area measurements from LC-MS analyses were log-transformed to normalize data distribution. A two-sample t-test was used to evaluate whether differences in PFOA serum concentrations existed between groups at different time points (Figure 1B). At baseline, low but detectible levels of PFOA were observed across all exposure groups but did not differ significantly across groups (*p* > 0.05). Following PFOA administration, the low- and high-dose groups exhibited significantly elevated concentrations compared to the control group at all post-exposure time points (*p* < 0.01 for days 7, 14, and 21). The greatest difference from control occurred at day 14, where the high-dose group showed an approximately 64% increase in serum PFOA relative to controls (Table S1). Concentrations remained consistently elevated in both PFOA-exposed groups throughout the study, with no statistically significant differences observed between the low- and high-dose groups after day 7.

**Figure 1.**
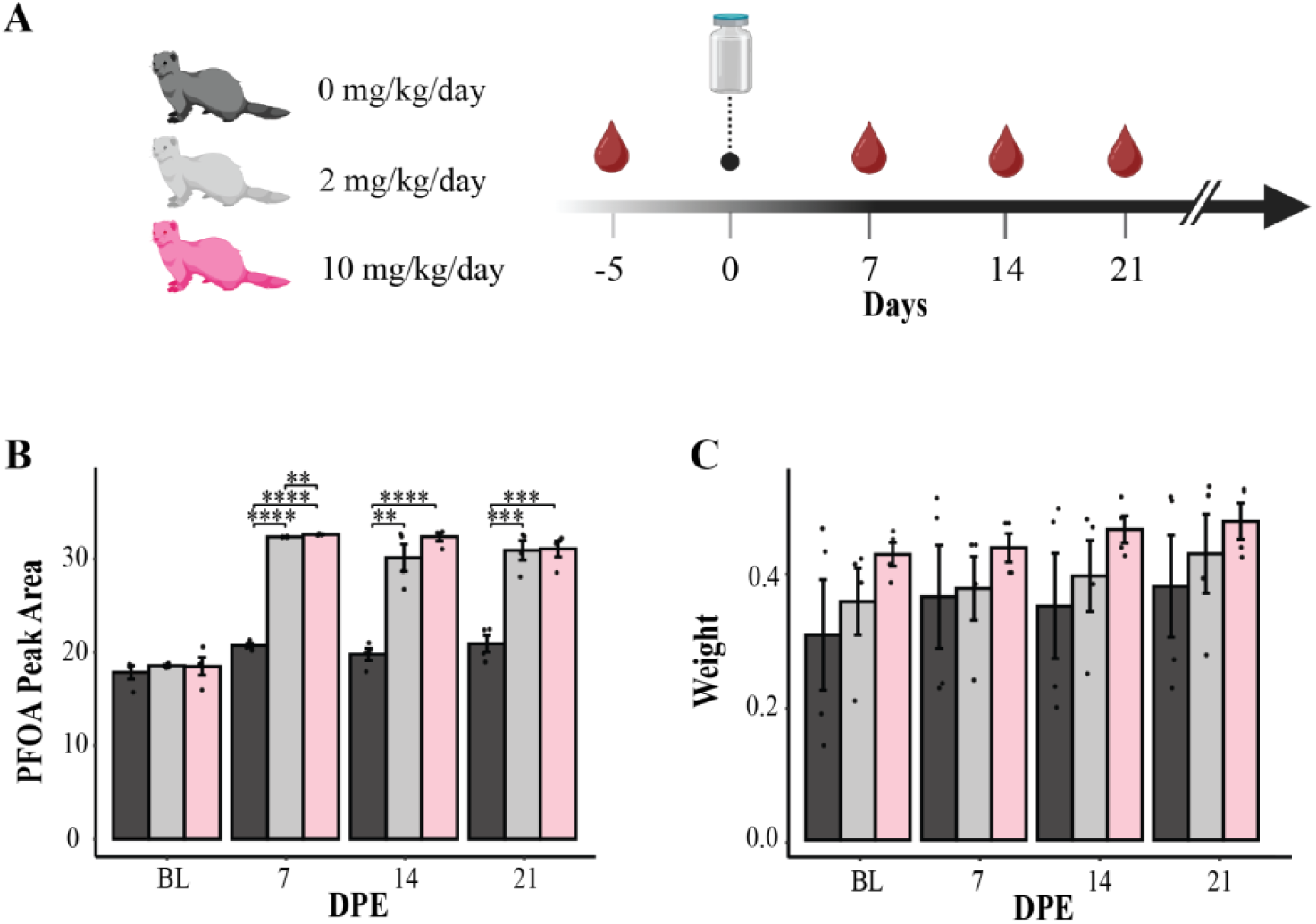
Sustained PFOA exposure in ferrets demonstrates successful dosing with no significant toxicity over 21 days (*n* = 4 per group). (A) Schematic of the experimental design depicting oral PFOA administration (0, 2, and 10 mg/kg/day) and the timeline for blood collection. *n* = 4 per group. (B) PFOA quantity in serum samples, represented by peak area, showing temporal changes of mean log-transformed peak area using LC-MS. Asterisks represent *p*-values measured by students t-test (\**p* < 0.05, \*\**p* < 0.01, \*\*\**p* < 0.001, \*\*\*\**p* < 0.0001) and error bars indicate standard deviation. BL, baseline; DPE, days post-exposure. (C) Log-transformed mean body weight trajectories (in kg) across exposure groups over the 21-day exposure period. Error bars indicate standard deviation. BL, baseline; DPE, days post-exposure.

Body weight is a commonly used marker of systemic toxicity ^[38]^, with decreased by weight reported in adult animals following high-dose PFOA exposure ^[9, 39]^. Daily body weight was monitored to evaluate potential PFOA-induced systemic toxicity (Figure 1C). Despite sustained PFOA exposure and increases in weight over time, there were no significant differences in body weight among exposure groups (*p* > 0.05). To further assess the relationship between PFOA exposure and body weight, linear regression analysis was performed on peak areas and body weight data. This analysis yielded a weak correlation coefficient (*r* = 0.228) and a non-significant p-value (*p* = 0.062), indicating that the effect of PFOA exposure on weight variance was minor (Figure S1). These findings confirm successful PFOA administration in a ferret model and sustained exposure while demonstrating minimal effects on body weight and thus, systemic toxicity throughout the 21-day exposure period.

**Table 1. Renal and liver chemistry profiles completed -7 and -1 days prior to challenge** Mean values for each analyte are shown for control ferrets at day -7 and PFOA-exposed ferrets at both day -7 and day -1. Data were analyzed using the Mann-Whitney U test to assess group differences. Asterisks indicate analytes that were significantly different between control (-7DPC only) and the PFOA-exposed groups at either time point with a false discovery rate (FDR) *q*-value < 0.05.

**Table 1.**
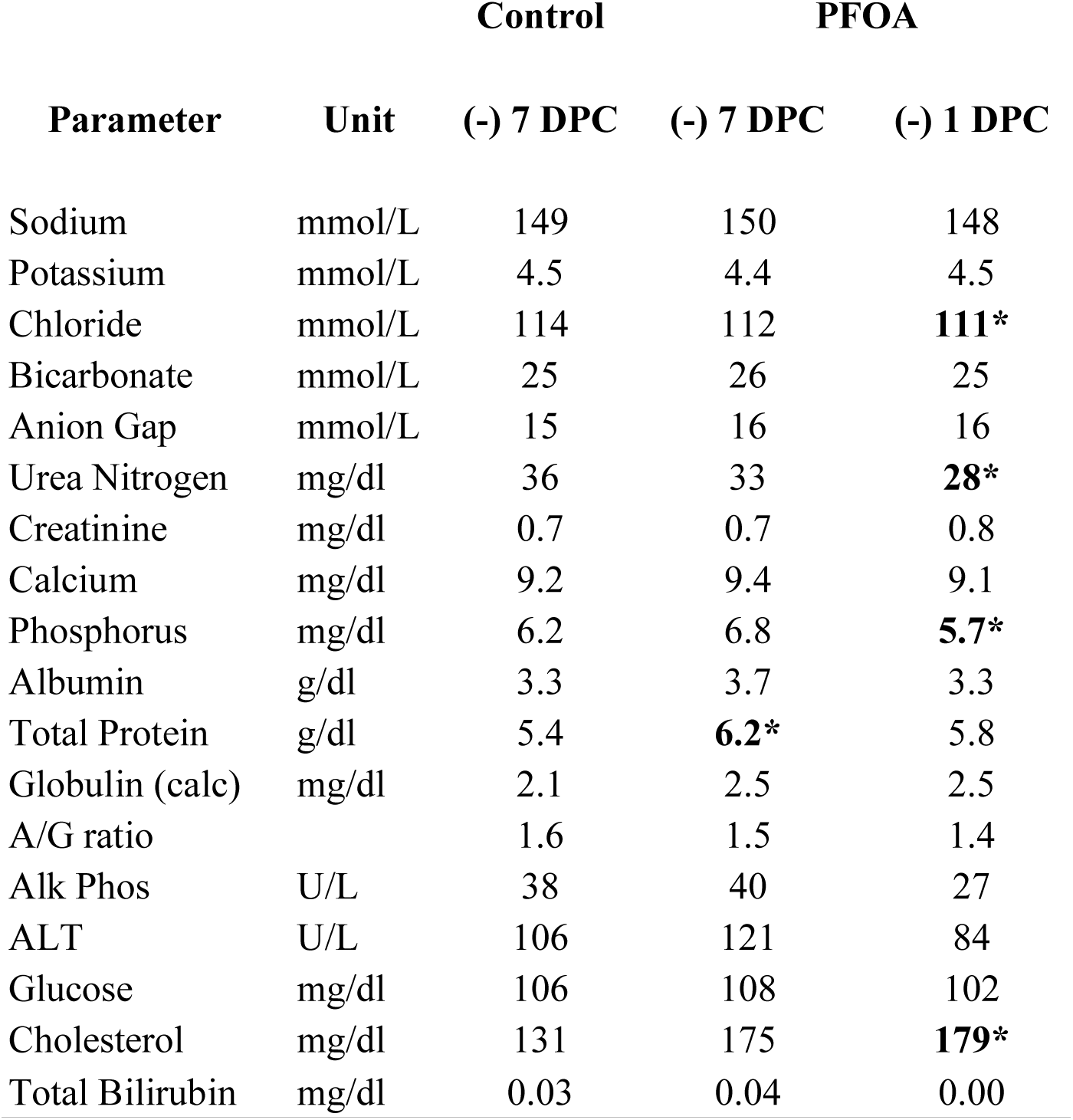
Serum chemistry profiles. completed -7 and -1 days prior to challenge. ALT: Alanine transaminase; Alk Phos: Alkaline phosphatase

### Results Section 2: PFOA Exposure Alters Blood Chemistry and Metabolic Profiles in Ferrets

Whole blood and serum samples were analyzed to investigate the impact of PFOA exposure on metabolites and liver and renal blood chemistry analytes. Whole blood samples were collected during the exposure phase of the study, one week (-7 days) and one day (-1 day) prior to SARS-2 challenge (Figure 2A). Sixteen ferrets were equally divided into control and PFOA-exposed (10 mg/kg/day) groups and administered their respective PFOA doses through feed for one week prior to SARS-2 infection. At day 0, all ferrets were intranasally inoculated with 1.0 ml PBS containing 10^6^ plaque forming units (PFU) SARS-2 (USA-WA1/2020, BEI Resources, NR-52281; SARS-2). This represents our “challenge” phase. Ferrets were monitored daily, and nasal washes, body weight, and temperature were collected every other day. Four days after the intranasal inoculation, a subset of animals (*n* = 3 per group) were humanely euthanized for the collection of early viral, immunological, and histopathological endpoints. At 13 days post-challenge (DPC), three control ferrets and two exposed ferrets were humanely euthanized. The remaining ferrets were humanely euthanized at 14 DPC. Tissue and blood samples were collected at each necropsy time point.

**Figure 2.**
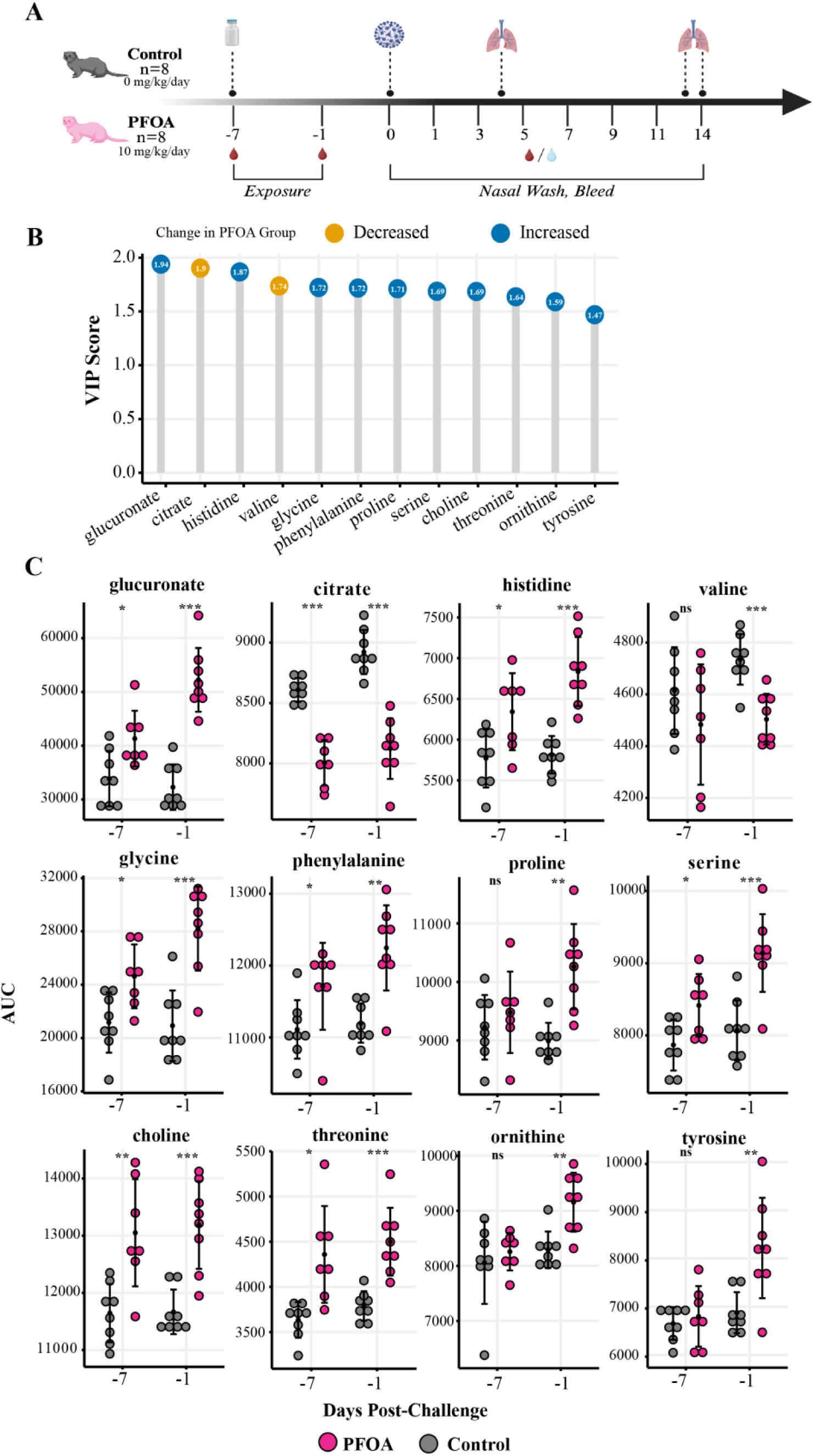
Temporal analysis of PFOA-induced changes in blood chemistry and metabolome prior to SARS-2 challenge. (A) Schematic representation of the experimental timeline, showing PFOA exposure (10 mg/kg/day), sample collection points, and SARS-2 viral challenge. (B) Variable importance in projection (VIP) scores for the top 12 metabolites identified by OPLS-DA (R^2^Y = 0.89, R^2^X = 0.59, Q^2^Y = 0.88). Metabolites that decreased in PFOA-exposed ferrets after one week of daily dosing are shown in yellow, while those that increased are shown in blue. (C) Plots of significantly altered metabolites identified at day -7 and day -1. Individual variability and group differences are depicted, with statistical significance denoted by adjusted *p*-value with false discovery rate correction (\**p* < 0.05, \*\**p* < 0.01, \*\*\**p* < 0.001, \*\*\*\**p* < 0.0001). n.s., no significance.

To investigate if PFOA exposure would affect metabolites in this model, whole blood samples from -7 and -1 DPC were processed for serum and used for untargeted metabolomic profiling by NMR. Significant features were collected using orthogonal partial least squares discriminant analysis (OPLS-DA) (R^2^Y = 0.89, R^2^X = 0.59, Q^2^Y = 0.88) and partial least squares discriminant analysis (PLS-DA; R²Y = 0.98, R²X = 0.39, Q² = 0.88) on the samples collected -1 DPC. Initial exploratory analysis and visualization of the data indicated less variance between the exposure groups at -7 DCP in comparison to -1 DPC (Figure S2). Based on variable importance projection (VIP) scores from the OPLS-DA and PLS-DA models, 85 spectral features corresponding to unique chemical shifts were prioritized for further analysis. These features were then annotated using COLMARm ^[40]^, STOCSY ^[41]^, and STORM ^[42]^ to identify corresponding metabolites. Metabolites were assigned a confidence score from 1 to 5, with 5 as the highest confidence score. The scores were defined as follows: 1) putatively characterized compound classes or annotated compounds, 2) matches from 1D NMR to the literature and/or other database libraries, 3) matched to HSQC, and/or STOCSY/TOCSY 4) matched to HSQC and validated by HSQC-TOCSY, validated by spiking the authentic compound into the sample 5) validated by spiking the authentic compound into the sample. Only metabolites with a score ≥3 were retained for downstream statistical analysis (Table S2). For each retained metabolite, a representative “driver peak” was selected.

Among these, glucuronate, citrate, and histidine exhibited the highest VIP scores (VIP > 1.87), indicating a higher contribution to the model (Figure 2B). Welch’s t-test hypothesis testing was applied to each dosing group within the exposure phase and corrected for FDR by Benjamini-Hochberg procedure. Figure 2C shows that 11 out of 12 of the metabolites increased in significance from -7 to -1 DPC.

To further investigate the effects of PFOA exposure in the ferret model, renal and liver blood chemistry analytes were assessed using serum (Table 1). A total of 18 analytes were measured from control ferrets at -7 DPC and from PFOA-exposed ferrets at both -7 DPC and -1 DPC. Statistical comparisons were performed using the Mann-Whitney U test, and significance was determined using FDR *q*-values (Table S3). Among the analytes, total protein was significantly elevated (*q* < 0.05) in PFOA-exposed ferrets at -7 DPC. By -1 DPC, chloride, urea nitrogen, phosphorus, and cholesterol levels were significantly different (*q* < 0.05) in the PFOA-exposed group compared to baseline control. Pearson correlation analysis of renal and liver analytes with serum metabolites indicates strong positive correlations between ornithine and creatinine, and between cholesterol with serine and glycine (Figure S3). Collectively, these findings indicate that PFOA exposure induces measurable and progressive disruptions in blood chemistry and serum metabolites.

### Results Section 3: Longitudinal Analysis Identifies Metabolic Alterations Associated with SARS-2 Infection and PFOA Exposure

On day 0 (0 DPC) of the infection phase, all ferrets were intranasally inoculated with SARS-2 (Figure 2A). Whole blood was collected every other day and processed into serum for NMR metabolomics analysis. The primary objective of the infection phase was to identify metabolites that changed significantly over time in response to SARS-2 infection while distinguishing between dosing groups. We aimed to identify metabolites consistently perturbed by PFOA exposure. To achieve these goals and account for the nonlinear longitudinal nature of the data, we employed a Generalized Additive Mixed Model (GAMM) to identify metabolites with significant associations to time (disease progression), dosing group, and the combined interaction of both. This analysis revealed 160 spectral features significantly associated with either the exposure (dose) effect, time effect, or both. These features were subsequently annotated and scored using database matching. From this set, 15 metabolites were identified with high confidence (Figure 3A).

**Figure 3.**
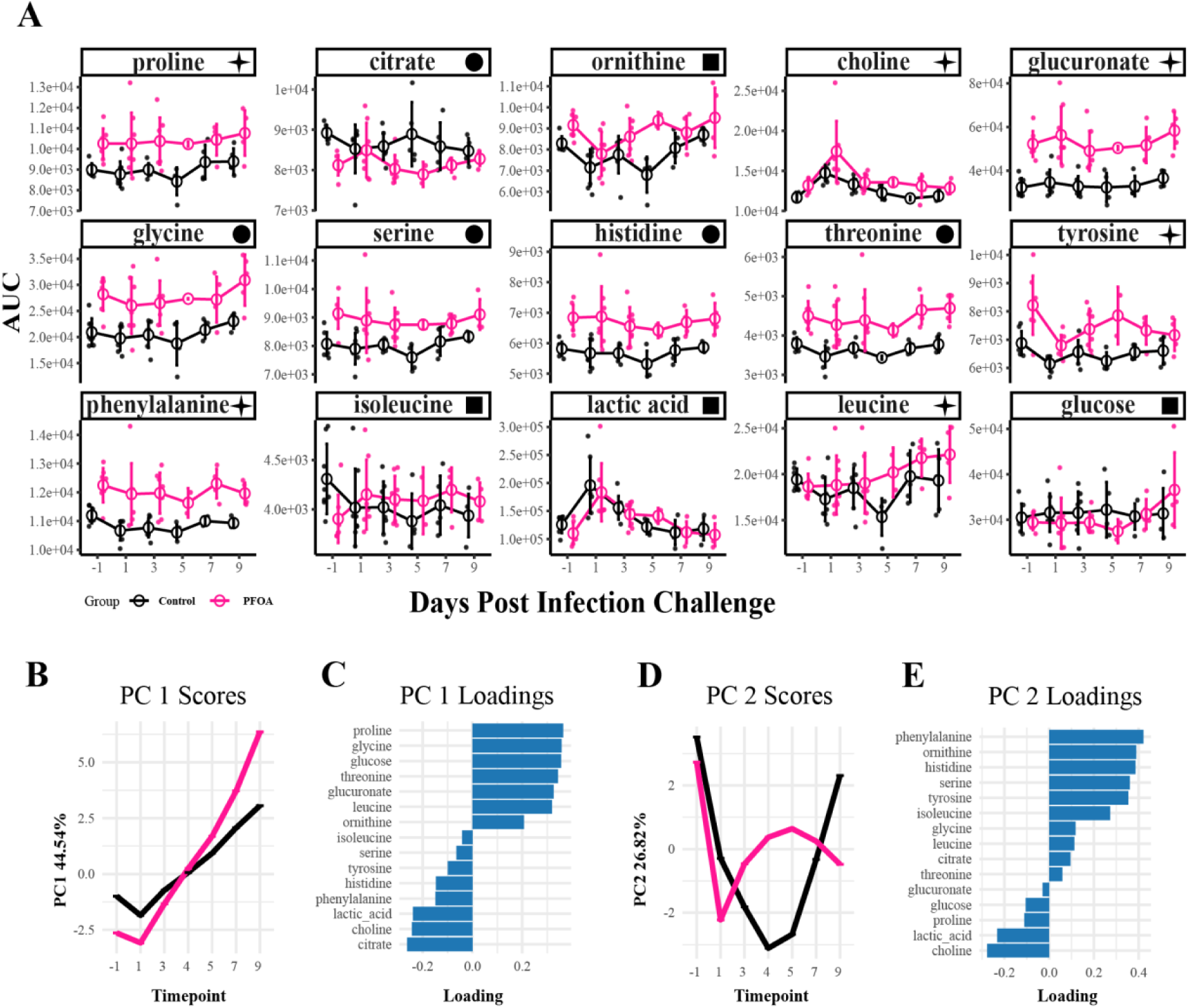
Metabolites with Significant Dose-Dependent and Temporal Changes During the Infection Phase. (A) Line plots of serum metabolites identified using generalized additive mixed models (GAMM), showing significant dose-dependent and time-dependent changes in response to SARS-2 infection. ■ indicates an infection response, ✦ indicates infection and dose response, ● indicates dose response only. (B-E) Principal component analysis (PCA) of GAMM-fitted residuals summarizing major patterns of metabolite variation. (B) PC1 scores across timepoints capture 44.54% of the model variation. (C) PC1 loadings indicate the metabolites contributing most to PC1 scores. (D) PC2 scores across timepoints capture 26% of the model variation. (E) PC2 loadings indicate the metabolites contributing most to PC2 scores.

Eleven of the 12 significant metabolites in the PFOA analysis (Figure 2C) remained significant post-infection (Figure 3). Valine was not significant after infection; however, leucine, isoleucine, lactic acid, and glucose were significant. Glycine, serine, histidine, threonine, and citrate exhibited significant dose-dependent effects (*p* < 0.05), with PFOA-exposed ferrets consistently differing from the controls throughout the infection study (Table S4). Proline, leucine, tyrosine, and glucuronate showed both significant dose effects and time-dependent changes within the PFOA group. Phenylalanine displayed a dose-dependent effect and showed significant temporal changes only in control ferrets. Glucose and ornithine exhibited time-dependent changes in PFOA-exposed ferrets, while isoleucine changed over time only in controls. Lactic acid showed significant temporal variation in both groups without a dose effect, whereas choline demonstrated both dose- and time-dependent differences across groups.

Principal component analysis (PCA) was then applied to the smoothed and fitted residuals to reduce the dimensionality and summarize the major patterns. PC1 captured 44.54% of the variance and showed increases over time after a decrease at 1 DPC (Figure 3B). The PFOA group diverged from the control group after 3 DPC. The associated loadings for this component indicated that proline, glycine, glucose, citate, choline, and lactic acid contributed most to the model variance (Figure 3C). The temporal trends observed in PC1 indicate a direct response to the infection challenge. PC2 explained 26% of the variance and captures more variable trends between the two groups (Figure 3D). The associated loadings showed phenylalanine, ornithine, and histidine have the highest contributions to the variation (Figure 3E). Although PC2 captured less variance in the model, it revealed temporal trends by group, indicating differences in responses to the infection challenge over time. Collectively, these results reveal that: 1) PFOA-induced metabolic changes persisted across the infection phase for select metabolites, 2) specific metabolites exhibited distinct temporal patterns in response to SARS-2 infection, and 3) the metabolic effects of PFOA exposure varied in magnitude and temporal progression depending on the specific metabolite.

### Results Section 4: PFOA-Exposed Ferrets Exhibit a More Severe Immune Response to SARS-2 Infection than Control Ferrets

To evaluate the extent of SARS-2 infection and potential differences in disease progression between PFOA-exposed and control ferrets, nasal wash samples were collected every other day for viral titer analysis by plaque assay. PFOA-exposed ferrets reached peak infection earlier, on day 3 post-challenge, whereas control ferrets reached peak infection on day 5 post-challenge (Figure 4A). Mixed-effects modeling of longitudinal viral titers, with Sidak’s multiple-comparisons post-hoc method ^[43]^, identified a significant difference between PFOA-exposed and control ferrets at 3 DPC (*p* = 0.0163). In addition to reaching peak infection sooner, PFOA-exposed ferrets exhibited a higher mean peak viral titers than controls (raw means 1200.6 PFU vs 540.0 PFU, respectively). These findings suggest that PFOA-exposed groups had accelerated viral replication and an increased viral burden, thereby exacerbated SARS-2 infection.

**Figure 4.**
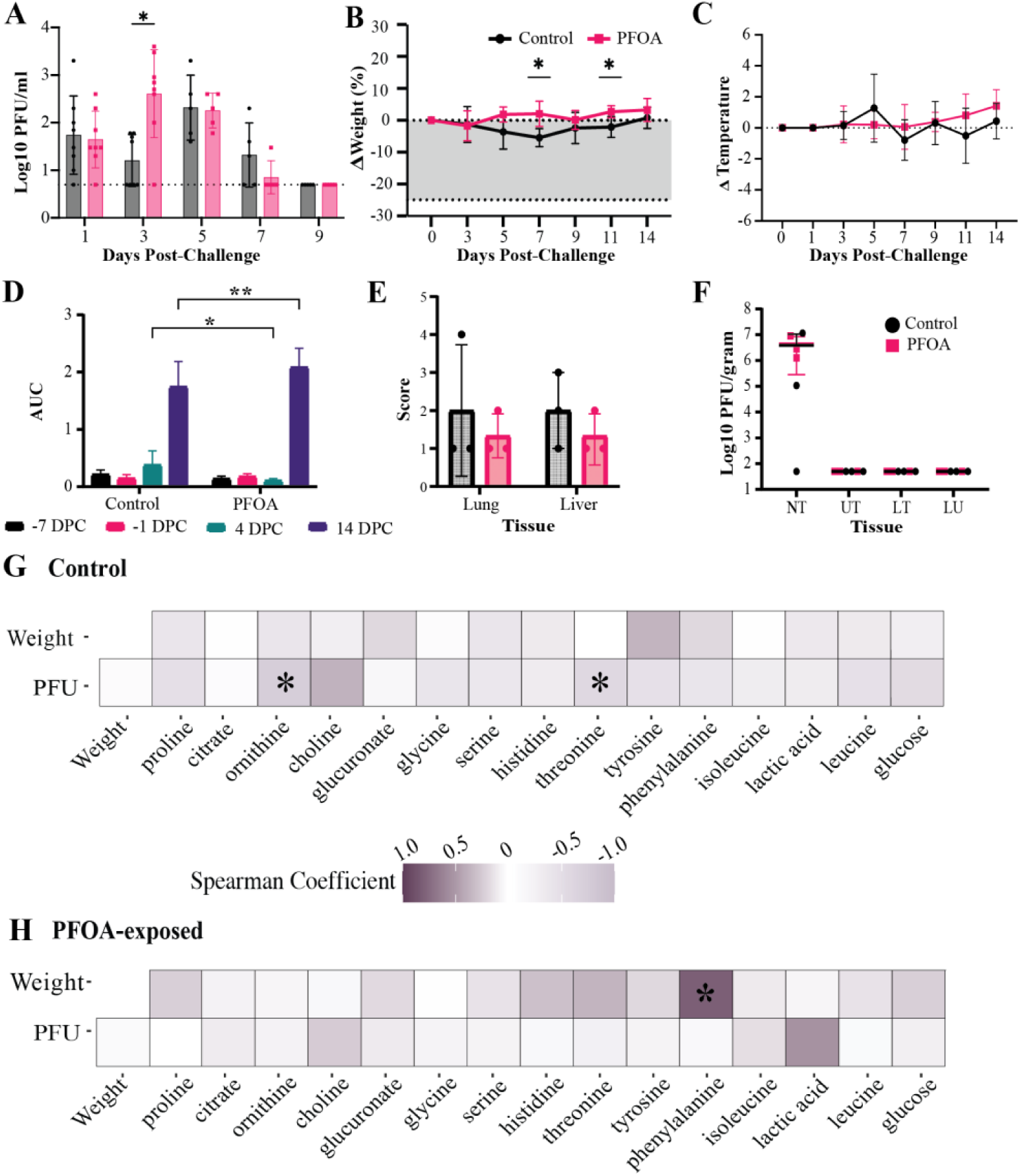
Infection and Immune Response to SARS-2 in Ferrets. (A) Plaque assay quantifying viral titers in ferret nasal wash samples over time, grouped by dosing group (*n* = 8 for days 1,3 and *n*=5 for days 5-9). The x-axis represents days post-SARS-2 challenge (DPC). (\**p* < 0.05) (B) Percentage weight changes following SARS-2 challenge (*n*=8 for days 0, 3 and *n*=5 for days 5-9). The thick dotted line indicates the 25% weight loss humane endpoint. The x-axis represents DPC. Asterisks denote statistically significant differences between Control and PFOA-exposed ferrets at individual time points based on mixed-effects modeling with Sidak’s multiple-comparisons correction. Within-group temporal comparisons are reported in Table S5. (\**p* < 0.05) (C) Percent ferret temperature changes (°F) recorded following SARS-2 challenge. The x-axis represents DPC. (*n*=8 for days 1, 3 and *n*=5 for days 5-9) (D) Ferret anti-SARS2 receptor binding domain RBD serum IgG levels shown as peak Area under the Curve (AUC), grouped by time points: day -7, day -1 day 4 and day 14. Statistical significance determined by two-way ANOVA with Dunnett’s multiple comparisons test. (E) Histopathological scoring of liver and lung tissues collected at 4 DPC. Bars represent group means (*n* = 2). Liver scores reflect the severity of hepatitis, while lung scores include assessments of interstitial pneumonia and perivascular leukocyte cuffing. No statistically significant differences were observed between groups. (F) Viral titers in tissues collected from nasal turbinates (NT), upper trachea (UT), lower trachea (LT), and lung (LU), shown as log_10_ plaque-forming units (PFU) per gram of tissue. (G) Spearman correlation heatmap for control ferrets, showing relationships between weight, viral titers (PFU), and metabolites. Asterisks indicate strong correlations (r > 0.5). (H) Spearman correlation heatmap for PFOA-exposed ferrets, showing relationships between weight, viral titers (PFU), and metabolites. Asterisks indicate strong correlations (r > 0.5).

Weight monitoring throughout the study revealed contrasting trends between the two groups. Control ferrets exhibited a gradual decrease in body weight following infection, reaching a maximum mean loss of 5.45% at 7 DPC (Figure 4B). In contrast, PFOA-exposed ferrets showed a slight but consistent increase in body weight over the same period, reaching a maximum mean gain of 3.21% at 14 DPC. Mixed-effects modeling of longitudinal weight change, with Sidak’s multiple comparisons, identified statistically significant differences between Control and PFOA-exposed ferrets at 7 DPC (*p* = 0.0101) and 11 DPC (*p* = 0.0313), as well as a significant overall effect of exposure (*p* = 0.0391) and a time-by-exposure interaction (*p* = 0.0349) (Table S5).

Body temperature fluctuations during the infection phase also differed between the groups (Figure 4C). Mixed-effects modeling of longitudinal temperature change, with Sidak’s multiple comparisons correction, revealed no statistically significant effects of time, exposure, or their interaction (all *p* > 0.05; Table S5). Control ferrets showed variable temperature changes, while PFOA-exposed ferrets experienced a gradual, though statistically insignificant, increase in body temperature.

Serological analysis of immunoglobulin G (IgG) levels provided further insights into the immune response. Control ferrets demonstrated an increase in serum IgG by day 4 post-infection, which was not observed in PFOA-exposed ferrets (Figure 4D). However, by day 14, IgG levels in the PFOA group had surpassed those of the control group, indicating a delayed yet full antibody response in PFOA-exposed ferrets. These results suggest that PFOA exposure may potentially delay the production of SARS-2-specific antibodies. To assess tissue-level pathological changes at 4 DPC, liver and lung samples were routinely processed and stained with hematoxylin and eosin (H&E). Histopathological evaluation was conducted by a board-certified veterinary pathologist blinded to exposure groups. Liver inflammation, or hepatitis, was scored based on the degree of periportal leukocyte infiltration, using the following scale: 1 = minimal, 2 = mild, 3 = moderate. Lung pathology was assessed by scoring the presence of interstitial pneumonia and perivascular leukocyte cuffing (Figure 4E). Overall, changes were mild and insignificant between groups. Additional tissues, including the kidney, heart, and spleen were also examined; however, they showed no significant histopathological findings (data not shown). Taken together, the lack of histopathological findings in examined organs supports that the administered PFOA doses were not systemically toxic.

To assess viral distribution across tissues, samples were collected 4 days post-infection from the nasal turbinate (NT), upper trachea (UT), middle trachea (MT), lower trachea (LT), cranial and caudal lung lobes, and the apex of the heart. Viral titers were quantified by plaque assay and expressed as PFU/gram of tissue. Analysis revealed detectable viral titers exclusively in the NT, with no significant differences in NT viral load between PFOA-exposed and control groups (*q* > 0.999) based on a Mann-Whitney test with FDR correction (Table S6).

Pearson correlation analysis was performed using plaque-forming unit (PFU) counts from nasal wash titer and body weight as disease progression indicators to explore potential relationships between metabolite levels and infection severity. In the control group, significant correlations were observed between nasal wash PFU titers and the metabolites ornithine (*r* = -.73, *p* < 0.05) and threonine (*r* = -.57, *p* < 0.05) (Figure 4G). No other significant correlations were identified in the control group. In contrast, PFOA-exposed ferrets exhibited no correlations between PFU titers and any metabolites. However, a significant correlation was observed between body weight and phenylalanine (*r* = -.81, *p* < 0.05) levels in the PFOA group (Figure 4H). These results together highlight the immune response and altered metabolic profiles in PFOA-exposed ferrets during SARS-2 infection.

## DISCUSSION

This study used an integrative clinical and metabolomics approach to investigate the effects of PFOA on COVID-19 disease severity. Through longitudinal sampling and serum metabolomics, we demonstrated that PFOA is detectable in serum and induces significant alterations in the metabolic profile within one week of daily oral exposure in a ferret model. PFOA-exposed ferrets experienced a more severe response to SARS-2 infection than controls, with distinct differences in metabolic trajectories over time.

### Ferret Exposure

PFOA’s immunotoxic effects have been well-documented in the context of influenza virus infection and vaccine response studies ^[13, 14]^; however, these immunotoxic properties remain elusive mechanistically, particularly regarding how metabolism contributes, and warrants investigation in a controlled setting. Ferrets offer a unique translational platform for this purpose, as they are widely used in infectious disease research due to their physiological similarity to humans, particularly in respiratory tract anatomy and immune responses ^[32]^. Ferrets are naturally susceptible to human respiratory pathogens, including SARS-2 ^[44–46]^, and they reproduce clinical and pathological features of human respiratory infections ^[44, 45]^. Their longer lifespan relative to rodents and ability to model disease severity over time make them a valuable system for studying both viral pathogenesis and environmental co-exposures ^[31, 47]^.

Despite their growing use in respiratory research, ferrets have not been commonly used to model PFAS exposure. In this study, low (2 mg/kg/day) and high (10 mg/kg/day) doses of PFOA were administered with mixed feed, based on established mouse model exposure models, to determine the presence or absence of systemic toxicity and dose selection for subsequent analysis ^[48]^.

PFOA was detectable in serum by LC/MS, and by days 14 and 21, serum PFOA levels plateaued, with no statistically significant differences between the low and high exposure groups. This plateau effect is consistent with observations in other animal models where repeated dosing leads to a rapid approach to steady-state serum concentrations despite PFOA’s long half-life. In primates, Butenhoff et al. reported that serum PFOA reached steady state within 4-6 weeks of daily oral dosing ^[49]^. Similarly, pharmacokinetic modeling and experimental data in rodents demonstrate that steady state can be achieved within days to weeks of repeated exposure ^[50, 51]^.

Regulatory reviews, including the Food Standards Australia New Zealand (FSANZ) critique, also highlight that steady-state serum levels are a common feature of chronic PFOA exposure ^[52]^. Our findings are consistent with observations in other species and support the relevance of the ferret model for evaluating the effects of PFOA exposure. Studies incorporating urine and tissue metabolomics, particularly from the liver and kidney, would strengthen the understanding of PFOA’s biodistribution and accumulation in ferrets. Weight is often used as a marker of systemic toxicity. We observed a non-significant increase in body weight across dosing groups over time.

### Blood chemistry in response to PFOA exposure

Once PFOA enters the body, it preferentially binds to proteins and accumulates in tissues with high protein content, particularly the kidney, liver, and serum, where it has been consistently detected ^[53–56]^. These bioaccumulations have been linked to a range of toxicities in human and animal models, including hepatotoxicity and immunotoxicity ^[14]^, and are known to interfere with both liver and kidney health and metabolism ^[14, 57–59]^. Kidney diseases have been linked to PFOA exposure in humans, with epidemiological studies showing associations with chronic kidney disease and renal cell carcinoma ^[60, 61]^. However, these findings have not been consistently reported in animal models, where most toxicity studies report little to no explicit kidney damage following PFOA exposure ^[62, 63]^. Due to the liver’s high protein content and the elevated expression of peroxisome proliferator-activated receptor alpha (PPARα), which is involved in regulating genes related to cell proliferation, inflammation, lipid modulation, and glucose homeostasis, the liver is one of the primary organs for PFOA accumulation in animals ^[9, 64–66]^.

The accumulation of PFOA in the liver can disrupt liver metabolism and cause dysfunction ^[67, 68]^. To assess early toxicity, we measured liver and kidney-associated serum analytes in controls and PFOA-exposed ferrets at days -7 and -1 relative to SARS-2 challenge. Of the 18 analytes assessed, chloride, phosphorus, BUN, total protein, and cholesterol were significantly altered in PFOA-exposed animals compared to baseline controls. We also observed a non-significant decrease in serum alkaline phosphatase (ALP) and alanine transaminase (ALT).

In general, increases in serum blood urea nitrogen (BUN) and creatinine are considered hallmark indicators of kidney dysfunction; however, there have been few consistent correlations found between PFOA exposure and elevated BUN or creatinine ^[63, 69, 70]^. In this study, after one week of daily PFOA exposure, we observed no clear signs of kidney damage in ferrets based on BUN and creatinine values. Serum creatinine remained within normal limits (WNL) for ferret reference ranges ^[71, 72]^. While observed values appear to fall WNL based on some selected published literature, there is considerable variability depending on biological factors such as age, sex, and reproductive status ^[72]^. BUN levels decreased significantly in PFOA-exposed ferrets, and were lower than normal ranges, however this value alone does not indicate clear kidney damage ^[71]^. Albumin and total protein, which are often affected in glomerular damage, also remained WNL. Although total protein showed a statistically significant increase, it was still WNL ^[71]^. PFOA is known to accumulate in the proximal tubule, where it interacts with key renal transporters that play roles in the secretion and reabsorption of electrolytes and organic anions ^[10, 73–75]^. While still WNL, both serum chloride and phosphorus were significantly decreased in the PFOA-exposed group. Histological analysis did not show severe signs of damage to the tissues/renal tubules between exposure groups. Taken together, these findings do not conclusively indicate early signs of kidney dysfunction and decline, and further studies are required to validate this interpretation.

In our study, one of the clearest associations between PFOA and liver metabolism was a significant increase in serum cholesterol following PFOA exposure. This elevation mirrors observations in human epidemiological studies, where elevated serum PFAS concentrations were consistently associated with increased total cholesterol ^[76–78]^. Unlike human studies, PFOA exposure induces hypocholesterolemia in rodent models ^[79]^. Our ferret model showing elevated serum cholesterol may better reflect the human response profile and is relevant for translational toxicology. However, there are potential confounding factors that may influence these findings. Analytes such as cholesterol are susceptible to pre-analytical variability, including sample handling, hemolysis, lipemia, and the fasting status of the animals ^[71]^. A key limitation of the study is the inability to reliably fast the animals, which may have contributed to variability in the parameters. ALP, a non-specific enzyme that can be present in a variety of tissues including the liver, intestines, bone, and kidneys, showed a non-significant decrease among groups. A reduction in serum ALP activity in ferrets is typically not considered clinically significant in isolation, particularly in the absence of other biochemical abnormalities or clinical signs. Unlike elevated ALP, which may be associated with hepatobiliary or skeletal pathology, low ALP activity is infrequently of pathological relevance. Elevated alanine transaminase (ALT) is a biomarker of hepatocellular damage and is a valuable liver parameter in ferrets ^[80, 81]^. ALT levels in PFOA-exposed ferrets decreased after 1 week of daily dosing but remained within reference levels. Some epidemiological studies have reported positive associations between PFOA and ALT ^[82–84]^; however, other studies have found no consistent relationship ^[85, 86]^. To our knowledge, there is no literature indicating a decrease in ALT after PFOA exposure, and in general, decreased ALT is not considered pathologically relevant.

Overall, given the relatively short duration of the study, it is possible that insufficient time elapsed for clear structural or functional renal and hepatic damage to develop or be biochemically detectible. Additionally, urinalysis and tissue analysis would help assess the effects more clearly.

### PFOA effects on metabolites and immune response

We examined whether short-term daily exposure to PFOA alters the host metabolic and immune response to SARS-2 infection in ferrets. We measured serum metabolites by NMR in control and PFOA-exposed ferrets pre- and post-infection and identified early and sustained metabolic alterations in PFOA-exposed ferrets. The results demonstrate that while SARS-2 infection elicited a metabolic response in both groups, PFOA-exposed animals had a distinct altered metabolic profile prior to infection that persisted throughout the disease course. This persistence was supported by our GAMM-PCA model, where more metabolites showed significant group (exposure) effects than time (infection) effects. This suggests that the PFOA-induced changes potentially influence the severity of the infection.

Untargeted serum metabolomics at days -7 and -1 relative to infection challenge revealed that group separation emerged over the course of the 1-week dosing period. The earlier time point at day -7 showed less variance between groups, suggesting the metabolic alterations emerged or intensified over the course of the 1-week dosing period. Using the OPLS-DA model VIP scores to select metabolites of interest, we confidently identified 12 metabolites contributing strongly to the discrimination of groups. Of these, there were multiple amino acids, metabolites linked to hepatic detoxification, and metabolites primarily associated with energy metabolism. Our findings are consistent with previous reports of PFOA-induced perturbations in amino acid homeostasis and nitrogen balance ^[12, 87, 88]^ and mitochondrial and TCA function ^[88, 89]^. Several of the metabolites retained the alterations following infection.

Following SARS-2 challenge, both groups exhibited characteristics of an immune response. Lactic acid levels spiked at day-1 post-infection in all animals, consistent with the glycolytic burst required for rapid effector cell activation and pro-inflammatory responses ^[90, 91]^. Citrate levels diverged by exposure groups. Control ferrets showed an increase in citrate, peaking at day 5, consistent with the peak infection day. Citrate is an abundant metabolite active in multiple pathways, but it has a demonstrated role in histone acetylation and anti-inflammatory immune regulation ^[92–94]^ . PFOA-exposed ferrets exhibited persistent decreased citrate from pre-infection, through the challenge phase. This decrease may suggest disrupted mitochondrial and TCA cycle function, known targets of PFOA toxicity. The difference in serum citrate bioavailability between groups may reflect impaired metabolic flexibility during the transition from pro- to anti-inflammatory responses, as they have different energetic needs. More targeted approaches are required to fully elucidate this response.

Ornithine, proline, and histidine were elevated in PFOA-exposed ferrets before and after the infection. In COVID-19 patient cohorts, higher ornithine levels have been associated with more severe disease cases and cytokine storm ^[95, 96]^. Alterations in histidine-related metabolism have been observed in severe COVID-19 cases, but these alterations are not consistent across studies ^[97, 98]^. Proline metabolism has been linked to proinflammatory immune response ^[99, 100]^ and was elevated in PFOA ferrets throughout our study. Exposure to PFAS, including PFOA, has been shown to induce heightened inflammatory responsiveness and alterations in amino acid metabolism ^[12, 101, 102]^. Within this context, the sustained elevation of proline observed here may reflect PFOA-driven dysregulation rather than a SARS-2-associated response, as prior studies of SARS-2 infection have reported proline to be either inversely associated with disease severity or unchanged ^[96–98, 103]^. Phenylalanine has been indicated as a potential biomarker of disease severity in COVID-19 patients ^[97, 98, 104, 105]^. In our study, phenylalanine in PFOA-exposed ferrets was consistently higher than control values pre- and post-infection. Tyrosine in our ferret model dropped at day 1 post-infection in PFOA-exposed ferrets to near control levels before rising again. Outside of this, tyrosine remained higher in PFOA-exposed ferrets pre- and post-infection. Tyrosine has been shown to have variable associations with COVID-19 severe cases; with some studies indicating positive associations ^[98]^ and others inverse ^[97]^.

Leucine, isoleucine, serine, glycine, and threonine all were all elevated pre-infection and exhibited dose effects post-infection. These amino acids are integral to biosynthesis, redox balance, and immune cell proliferation. Literature on their role in COVID-19 severity is inconsistent with reports of both positive and negative associations ^[96–98, 103, 106]^. Choline and glucuronate were persistently elevated pre- and post-infection. While their direct role in antiviral burden is unclear, their stable elevation supports the interpretation of ongoing metabolic effects in response to PFOA dosage. The metabolite alterations observed align with the immune phenotypes measured in this study. PFOA-exposed ferrets reached higher peak viral titers earlier (day 3) than controls (day 5) and exhibited a delayed IgG antibody response. This delayed response is consistent with what we expected from influenza studies of PFOA immunotoxicity ^[14, 107]^. The suppression of altered citrate and amino acid profiles may reflect a metabolic environment less favorable for efficient immune response. Notably, in the control groups, ornithine and threonine were positively correlated with viral titers. No such correlations were observed in the PFOA group.

### Limitations and Future directions

Some metabolite behaviors in our study diverge from trends reported in human COVID-19 cohorts and PFOA studies, which may reflect a species-specific response or the controlled nature of our model. It is important to note that metabolite abundances alone cannot distinguish between increased synthesis or decreased utilization. Elevated metabolites could result from either enhanced production or reduced incorporation into downstream biosynthetic pathways.

Targeted pathway and metabolic flux analysis will be essential to resolve these mechanistic uncertainties.

## Conclusion

In summary, this work demonstrates that a short-term PFOA exposure produces distinct and persistent alterations in serum metabolite profiles. Many of the metabolites have immunometabolism functions relevant to SARS-2 pathogenesis. The fact that these alterations were established prior to infection and were sustained through disease resolution suggests that environmental exposures can preconfigure host metabolic states in a way that shapes infection trajectories. These findings underscore the need to consider environmental toxicant exposure history in infectious disease models and highlight metabolic pathways that warrant deeper investigation for their potential role in susceptibility and recovery.

## RESOURCE AVAILABILITY

### Lead Contact

Further information and request for resources and reagents can be directed to the lead contact, Arthur Edison

### Data and code availability

- Metabolomics data have been deposited at the NIH Common Fund’s National Metabolomics Data Repository (NMDR) website, the Metabolomics Workbench, https://www.metabolomicsworkbench.org where it has been assigned Study ID ST004607. The data can be accessed directly via its Project doi: http://dx.doi.org/10.21228/M85G21.
- All original code has been deposited on GitHub and is publicly available at (https://github.com/edisonomics/PFAS_COVID)
- Any additional information required to reanalyze the data reported in this paper is available from the lead contact upon request

### Materials availability

This study did not generate new unique reagents

## ACKNOWLEDGMENTS

**Funding:** This study was supported by research grants from the National Institutes of Health (NIH): P42 ES027704, 3U2CES030167-03S2, 5R35GM148240-03.

## AUTHOR CONTRIBUTIONS

D.N.L., D.R.H., M.U., J.R.C, F.E.L., J.D., T.W., D.A.G., E.S.B., F.M.F., S.M.T., and A.S.E. conceptualized the study. A.S.E., S.M.T., E.S.B., F.M.F., and J.D. acquired funding. A.S.E. provided study supervision. D.N.L., M.U., and D.R.H., curated the data. D.R.H. completed administration of PFOA, viral inoculation, viral quantification, serology, serum collection, and formal analysis. C.J., K.S., and S.J. completed clinical pathology and histopathology. F.E.L. completed PFOA measurement. D.N.L completed the NMR metabolomics sample preparation, data acquisition, processing, analysis, all coding and implementation of code, and formal analysis. D.N.L and D.R.H prepared all visualizations and wrote the manuscript. All authors provided comments.

## DECLARATION OF GENERATIVE AI AND AI-ASSISTED TECHNOLOGIES IN THE WRITING PROCESS

During the preparation of this work, ChatGPT and Grammarly were used to polish grammar and spelling. After using these tools, we reviewed and edited the content as needed and take full responsibility for the content of the publication.

## SUPPLEMENTAL INFORMATION

Document S1. Figures S1-S3 and Tables S2 & S4

Table S1 PFOA and Weight analysis, related to Figure 1

Table S3 Blood chemistry analytes analysis, related to Table 1

Table S5 Infection study analysis, related to Figure 4

Table S6 Tissue titer analysis, related to Figure 4

## METHODS

### Administration of PFOA

Sixteen, nine-month-old, male fitch ferrets were received from Triple F Farms (Pennsylvania, USA) and acclimated in the UGA animal facilities. Prior to arrival, ferrets underwent sterilization and scent gland removal. Following a period of acclimation, ferrets were anesthetized for subcutaneous placement of IPTT300 thermal transponders (Bio Medic Data Systems, USA) and bled. Prior to infectious challenge, dosing and subsequent loading of PFOA was conducted in two phases.

In the pilot dosing phase, ferrets were divided into three groups (n=4) and administered perfluorooctanoic acid (PFOA) at doses of 0, 2, or 10 mg/kg/day (Sigma-Aldrich 171468, batch WXBD3549V). PFOA was dissolved in water and combined with Gerber Turkey baby food to a final volume of 5.0mL. Mock ferrets were provided baby food only. Under direct observation, ferrets were fed the mixture once daily for twenty-one days. Following this initial dose analysis, PFOA was suspended for five weeks.

For the loading portion of PFOA delivery, ferrets were divided into two groups (n=8) and administered 0 or 10mg/kg/day PFOA in feed for seven days prior to infectious challenge. Ferrets in the 10 mg/kg treatment group had previously received either 2 or 10 mg/kg PFOA during the pilot. Whole blood samples were collected weekly.

Clinical observations, including weight and temperature, were recorded every other day throughout the administration of PFOA. Additionally, whole blood samples were collected from the ferrets weekly as well as one day prior to infectious challenge.

All experimental procedures prior to infectious challenge were performed at the University of Georgia Biosciences Animal Facility. For SARS-CoV-2 challenge, animals were transferred to the University of Georgia Animal Health Research Center Animal Biosafety Level -3 (ABSL3) facility. Animals were housed solo or in pairs as study design permitted, with ad libitum access to food and water. All experiments were approved by the IACUC of the University of Georgia College of Veterinary Medicine. All procedures including nasal washes and blood collections were performed under short term isoflurane gas anesthesia. UGA AUP: A2020 03-016-Y1

### PFOA Measurement

To monitor the PFOA uptake during the dosing phase, blood samples were analyzed using a Thermo Fisher Scientific Q Exactive HF Orbitrap mass spectrometry coupled to a Vanquish UPLC. Chromatographic separation was performed at 0.4 mL/min with an Agilent C18 column (2.1 x 150 mm, 1.8 mm, 30°C column temperature) over 24 minutes starting with 90 % H_2_O, 5 mM ammonium acetate (A), and 10% methanol (B) held isocratic for 0.5 minutes, then increased to 30% B at 2 minutes, 95% B at 14 minutes and then 100% B at 14.5 minutes and held for 2 minutes before re-equilibration at 10% B for 7 minutes prior the next injection. 2 mL of sample was injected per run. The sample queue was randomized with injection blanks included to monitor for sample carryover. All samples were analyzed by negative mode electrospray ionization (ESI). Full MS scans were performed at a specified resolution of 60,000 (m/z 200) from 150 to 2,000 m/z with an AGC target of 1e6 and a maximum IT of 200 ms.

### Virus

SARS-CoV-2 virus (USA-WA1/2020, BEI Resources, NR-52281; SARS-2) was propagated in Vero E6 cells cultured in Dulbecco’s modified Eagle medium (DMEM) containing 2% fetal bovine serum (FBS) and antibiotics at 37°C, 5% CO2. Infectious virus was titered in Vero E6 cells by plaque forming unit (PFU) and focus forming unit (FFU) assays.

### Viral inoculation and sampling

Ferrets were inoculated intranasally with 1.0 mL PBS containing 10^6^ PFU SARS-2, distributed equally between nares. Following inoculation, animals were monitored daily for clinical signs, including nasal discharge, sneezing, diarrhea, lethargy, increased respiratory rate and effort and altered responses to external stimuli. Nasal washes, weight and body temperature were collected at least every other day. Blood was collected one day before and every other day after SARS-2 inoculation. Whole blood was processed for serum and peripheral blood mononuclear cells (PBMCs). Four days after challenge, a subset of three animals per group was humanely euthanized. Blood and tissues were processed for viral titers and downstream analysis. The remaining animals were humanely euthanized two weeks after SARS-2 inoculation. Tissues including the nasal turbinates, trachea and cranial left lung lobe were placed in either viral transport media (VTM) or Trizol for virology. Additionally, portions of the nasal turbinates, trachea, left cranial lung lobe, thymus, heart, liver, right kidney, spleen and select adipose tissues (inguinal and mesenteric) were frozen at -80°C. Untreated urine and serum were also stored at - 80°C. The remaining portions of the head, trachea, lung, thymus, heart, liver, spleen, and the left kidney were stored in a 10% neutral buffered formalin solution for histology.

### Viral quantification

Nasal wash samples were serially diluted in DMEM, transferred onto cell culture plates containing confluent Vero E6 cells and incubated for 1 hour at 37°C, 5% CO_2_. Overlay media (OptiMEM containing 0.8% methylcellulose, 2% FBS and 1x A/A) was applied and plates incubated for 72 h at 37°C, 5% CO_2_. Overlays were decanted, plates washed, fixed and crystal violet developed. Plaque forming units were then manually calculated and quantified.

### Serology

To detect ferret serum IgG, medium-binding 96-well ELISA microplates (Greiner Bio-One, 655001) were coated with 20 μg of either the full length SARS-CoV-2 receptor binding domain (RBD) in sterile 1x phosphate buffered saline (Corning, 21-040-CV) overnight at 4°C. Plates were washed 3x with 300 µL of 0.05% PBS-T using an automated plate washer (BioTek 405 TS Washer). All washes were performed using the same technique. Wells were blocked with 200 µL Blocking Buffer (3% non-fat dry milk in 0.05% PBS-T), incubated 2 hours at room temperature, then washed. Heat inactivated serum was serially diluted in dilution buffer (1% non-fat dry milk in 0.05% PBS-T),100 μl added to the appropriate wells, and incubated 2h at room temperature followed by a wash step. Goat anti-ferret IgG HRP-conjugated antibody (Bethyl Laboratories, A140-108P) diluted 1:10,000 in dilution buffer was added at 100 μl per well and incubated for 1h at room temperature. Plates were washed with 0.05% PBS-T, then H_2_0 and tapped dry to remove residual solutions. To develop, 50 μl TMB Peroxidase Substrate (HRP) (Vector Laboratories, SK-4400) was added to each well. Following a 10 min development period, 50 μl 1 N H_2_SO_4_ was added to stop the reaction, and plates were read at absorbance of 450 nm (BioTek, Cytation7 machine). The background signal was determined by calculating the average absorbance value plus three times the standard deviation (SD) of absorbance values obtained from capture protein-coated wells that were treated with the goat-anti ferret IgG-HRP antibody.

### Clinical Pathology

Whole blood samples were collected weekly. Complete blood cell counts, leukograms, renal and liver profiles were processed through the University of Georgia College of Veterinary Medicine Veterinary Teaching Hospital (VTH) Clinical Pathology Services. Manual differentials and morphological evaluation were completed by trained clinical pathology laboratory technicians at the University of Georgia.

### Histopathology

Following sectioning, processing occurred in a Sakura VIP6 tissue processor. Tissues were dehydrated in a series of graded ethanol baths, washed with xylene, and soaked in paraffin for infiltration, then embedded in appropriately sized molds, sectioned at 4 µm and mounted on charged slides. Sections were dried in a 60°C oven, processed through a series of xylene baths to deparaffinize them, leaving only the tissue on the slides. The sections were then rehydrated through a series of graded alcohol baths, rinsed, and stained with Hematoxylin and Eosin (H&E) on a Sakura Prisma Automated stain machine and covered by a Sakura Glas2 automated cover slipping machine.

Slides were examined by a board-certified veterinary pathologist blinded to the groups. Briefly, alveolar infiltrates and bronchiolitis were scored based on distribution, with 1 representing focal lesions, 2 multifocal, 3 coalescing, 4 affecting most of the section, and 5 diffuse. Perivascular cuffing by leukocytes was scored based on thickness, with 1 representing a single layer of leukocytes, 2 = 2 – 5 layers, 3 = 6 – 10 layers, and 4 > 10 layers. Interstitial pneumonia was scored based on the thickness of the alveolar septa, with 1 representing a single leukocyte thickness, 2 = 2 cells thick, 3 = 3 cells thick, 4 = 4 cells thick. Damage to lung tissue was scored using epithelial hypertrophy, epithelial hyperplasia, presence of pseudo squamous epithelium and epithelial necrosis of the bronchi and bronchioles, hypertrophic type 2 pneumocytes, hyperemia septa, alveolar emphysema and alveolar hemorrhages. Epithelium changes were assigned values based on severity as: 0 = absent, 1 = minimal, 2 = slight, 3= moderate, 4 = strong, and 5 = severe. Acute lung inflammation was scored based on bronchitis, bronchiolitis, peribronchitis, peribronchiolitis, interstitial infiltrate, alveolitis, vasculitis and perivasculitis. Values assigned to determinants of acute inflammation, except alveolitis, were assigned according to layers of leukocytes present in the tissue. Alveolitis was scored 1 to 5 based on distribution as described for lung damage.

### Statistical Analysis of clinical chemistry and immune response data

Mann-Whitney U tests were used to compare PFOA dosed samples at days -7 and -1 against control day -7 (considered a baseline measurement for this analysis) for each blood chemistry marker. Adjusted *p*-values were calculated by Benjamin-Hochberg false discover rate correction and values less than 0.05 were considered significant. Group comparisons of serum IgG levels and tissue histology were conducted using two-way ANOVA with Dunnett’s multiple comparisons test. Comparisons of nasal shedding, weight, and temperature were conducted using mixed-effects modeling with Sidak’s multiple-comparisons correction. Analysis was carried out using Graphpad Prism software.

### NMR Buffer Preparation

A phosphate buffer was prepared by dissolving anhydrous sodium dihydrogen phosphate (NaH₂PO₄) in deuterium oxide (D₂O). A 0.1 M solution of DSS-d₆ (4,4-dimethyl-4-silapentane-1-sulfonic acid, sodium salt) was added to the buffer. DSS is used as a chemical shift reference (0.0 ppm). The pH was adjusted to 7.0, and the final volume was corrected with D₂O. The prepared buffer was aliquoted and stored at 4°C until use. Detailed protocol can be accessed via its protocol doi: dx.doi.org/10.17504/protocols.io.8epv5rj6jg1b/v1.

### Serum Collection and Extraction

Whole blood was collected from each ferret using a 3.5 mL CAT Serum Separator Clot Activator Vacuette (Greiner Bio-One, Ref 45228P). Tubes were centrifuged at 1500 × *g* for 10 minutes at 4°C to separate the serum. The isolated serum was mixed in a 1:3 ratio with ice-cold methanol (Fisher, A456-4 Lot 173812TF), vortexed briefly, and stored at -80°C until metabolite extraction. Serum samples were thawed on ice and aliquoted (800 µL per sample). Samples were vortexed at 4°C for 20 minutes and centrifuged at 16,000 rcf for 30 minutes. The supernatant was transferred to a clean tube and dried using a speed vacuum concentrator (SC210A SpeedVacPlus, Thermo Savant, USA). Dried pellets were resuspended in 600 µL of the phosphate buffer described above, vortexed for 15 minutes until fully dissolved, then centrifuged briefly (5–10 seconds). The solution was transferred to 5 mm SampleJet NMR tubes (Bruker Biospin, Billerica, MA, USA) for spectral acquisition. Detailed protocol can be accessed via its protocol doi: dx.doi.org/10.17504/protocols.io.8epv5rj6jg1b/v1

### Quality Control and Assurance

All study serum samples were randomized prior to extraction protocol and running NMR experiments. Six NMR buffer blanks and two extraction blanks were added. Each rack contained three buffer blanks, and an extraction blank. Two external reference material samples were included. Reference material samples were created using the iterative batch averaging method (IBAT) of non-study ferret serum samples ^[110]^. 25 µL were taken out of each study sample for six internal pooled controls. Half of the internal controls were pre-challenge, and the other half post-challenge. 1D ^1^H NMR experiment was conducted on all samples, while 2D NMR was carried out on one pre-challenge and one post-challenge internal control. The NMR data for the internal pooled samples were used for metabolite annotation.

### NMR Data Acquisition and Processing

Spectra were acquired on an Avance III HD 600 MHz Bruker NMR spectrometer. The following NMR experiments were conducted: one-dimensional nuclear Overhauser effect pulse sequence with presaturation of water resonance (NOESYPR1D), PROJECTpr1D, 1D J-resolved spectroscopy (JRES), two-dimensional (2D) ^1^H–^13^C heteronuclear single quantum correlation (HSQC), 2D total correlation spectroscopy (TOCSY), 2D HSQC-TOCSY. Detailed data acquisition parameters for individual NMR experiments, raw data files, and processed data files can be found on Metabolomics Workbench ^[108]^ and accessed directly via its project doi: http://dx.doi.org/10.21228/M85G21.

PROJECTpr1D spectra were phase and baseline-corrected using MestReNova with an in-house processing script. Chemical shift referencing to DSS was confirmed using the Edison Lab Metabolomics Toolbox (https://github.com/edisonomics/metabolomics_toolbox). The ends of the NMR spectra (less than -0.5 ppm and greater than 10 ppm), water region (less than 4.70 ppm and greater than 4.85 ppm), and methanol region (less than 3.36 ppm and greater than 3.33 ppm) were removed from all samples. Alignment was carried out using constrained correlation optimized warping (CCOW) ^[111]^. All sample peaks were manually binned and quantified to reduce noise, yielding features (unique chemical shifts) for each sample by taking spectral area for integration in regions without overlap. Processing and manual binning scripts for this dataset can be found on GitHub (https://github.com/edisonomics/PFAS_COVID). Binned spectral data were normalized using probabilistic quotient normalization (PQN) and log-transformed in R.

### Feature Annotation

2D NMR spectra (HSQC, TOCSY, and HSQC-TOCSY) were processed using NMRPipe ^[112]^. Metabolites were identified using COLMARm database-matching ^[40]^, statistical total correlation spectroscopy (STOCSY) ^[41]^, and subset optimization by reference matching (STORM) ^[42]^.

Metabolites were assigned a confidence score from 1 to 5, with 5 as the highest confidence score. The scores were defined as follows: 1) putatively characterized compound classes or annotated compounds, 2) matches from 1D NMR to the literature and/or other database libraries such as BMRB ^[113]^ and HMDB ^[114]^, 3) matched to HSQC, and/or STOCSY/TOCSY 4) matched to HSQC and validated by HSQC-TOCSY, 5) validated by spiking the authentic compound into the sample. Only metabolites with a score ≥3 were retained for downstream statistical analysis. For each retained metabolite, a representative “driver peak” was selected.

### Feature selection and analysis

Exploratory analysis was conducted using Principal Component Analysis (PCA) in MATLAB and R to visualize clustering and detect outliers. Samples that were outliers in the full-resolution spectra or fell outside the 95% confidence ellipse in PCA were excluded.

Features were selected in both phases of the study. To select metabolites based on the effects of PFOA exposure before SARS-CoV-2 challenge (day -1), orthogonal partial least squares discriminant analysis (OPLS-DA; R²Y = 0.89, R²X = 0.59, Q² = 0.88) and partial least squares discriminant analysis (PLS-DA; R²Y = 0.98, R²X = 0.39, Q² = 0.88) models were applied using the *ropls* package in R. Model performance was evaluated using cross-validation by data splitting, with odd-numbered samples used for training and even-numbered samples reserved for testing. Variable Importance Projection (VIP) scores from each model were used to rank features. The overlapping top-ranked features across both models were retained for identification. After identification, adjusted *p*-values (obtained from Welches t-test followed by Benjamini-Hochberg false discovery rate correction) were calculated between groups at day -7 and -1 to observe the difference after 1 week of daily dosing.

After challenge, significant features were selected using generalized additive mixed models (GAMM) to account for the nonlinearity, unbalanced, and longitudinal nature of the data. The model included fixed effects for each group (PFOA vs control), random effects for each ferret, and smooth terms for time. Features that were significant for time, group, or time x group interaction were retained for identification as described above. After identification, fitted effects of the significant metabolites were summarized using PCA for dimensionality reduction.

After challenge, significant features were selected using generalized additive mixed models (GAMMs) to account for nonlinear time effects, repeated measures, and unbalanced sampling across subjects. For each feature we modeled fixed effects for treatment group, smooth functions of time specified for each group, and random intercepts for subjects to account for the within-ferret correlation across timepoints. Smooth terms were estimated using penalized regression splines with restricted maximum likelihood. For each feature, statistical significance was evaluated for: 1) the parametric group effect, 2) the time effect within the control group, 3) the time effect within the PFOA group. Features with *p*<0.05 for any of these terms were retained for identification and used for multivariate modeling.

After fitting GAMMs, we applied PCA to the GAMM effects matrix for the identified metabolites. This framework was conceptually adapted from repeated-measures ANOVA-simultaneous component analysis (RM-ASCA+) ^[115]^, which combines mixed modeling with principal component analysis to decompose structured longitudinal effects. PCA was applied to the effect matrix to summarize dominant multivariate response patterns. The scores represent the major longitudinal response patterns across samples. The loadings represent the contribution of each metabolite to a given response pattern. Metabolites with large absolute loadings contribute more strongly to the trends described by that principal component.

Correlation analysis between significant metabolites and clinical carriable (blood chemistry, nasal shedding, weight, and temperature) was performed using Spearman correlation. Features with an absolute correlation coefficient |r| > 0.6 were considered relevant.

## Notes

http://dx.doi.org/10.21228/M85G21

### Competing Interest Statement

The authors have declared no competing interest.

## References

1. Li, D. and S. Suh, Health risks of chemicals in consumer products: A review. Environ Int, 2019. 123: p. 580–587.

2. @CDCEnvironment. *Fast Facts: PFA*S in the U.S. Population | PFAS and Your Health | ATSDR. 2024 2024-11-07T09:43:25Z; Available from: https://www.atsdr.cdc.gov/pfas/data-research/facts-stats/index.html.

3. Cousins, I.T., et al., Why is high persistence alone a major cause of concern? Environmental Science: Processes & Impacts, 2019. 21(5): p. 781–792.

4. Buck, R.C., et al., Perfluoroalkyl and polyfluoroalkyl substances in the environment: Terminology, classification, and origins. Integrated Environmental Assessment and Management, 2011. 7(4): p. 513–541.

5. Zahm, S., et al., Carcinogenicity of perfluorooctanoic acid and perfluorooctanesulfonic acid. The Lancet Oncology, 2024. 25(1): p. 16–17.

6. Vierke, L., et al., Perfluorooctanoic acid (PFOA) — main concerns and regulatory developments in Europe from an environmental point of view. Environmental Sciences Europe, 2012. 24(1): p. 1–11.

7. Lau, C., Perfluorinated Compounds: An Overview, in Toxicological Effects of Perfluoroalkyl and Polyfluoroalkyl Substances, J.C. DeWitt, Editor. 2015, Springer International Publishing: Cham. p. 1–21.

8. Ahrens, L. and M. Bundschuh, Fate and effects of poly- and perfluoroalkyl substances in the aquatic environment: A review. Environmental Toxicology and Chemistry, 2014. 33(9): p. 1921–1929.

9. Jiang, Q., H. Gao, and L. Zhang, Metabolic Effects PFAS, in Toxicological Effects of Perfluoroalkyl and Polyfluoroalkyl Substances, J.C. DeWitt, Editor. 2015, Springer International Publishing: Cham. p. 177–201.

10. Kudo, N., Metabolism and Pharmacokinetics, in Toxicological Effects of Perfluoroalkyl and Polyfluoroalkyl Substances, J.C. DeWitt, Editor. 2015, Springer International Publishing: Cham. p. 151–175.

11. Chen, Z., et al., Dysregulated lipid and fatty acid metabolism link perfluoroalkyl substances exposure and impaired glucose metabolism in young adults. Environment International, 2020. 145: p. 106091.

12. Prince, N., et al., Plasma concentrations of per- and polyfluoroalkyl substances are associated with perturbations in lipid and amino acid metabolism. Chemosphere, 2023. 324: p. 138228.

13. Keil, D.E., Immunotoxicity of Perfluoroalkylated Compounds, in Toxicological Effects of Perfluoroalkyl and Polyfluoroalkyl Substances, J.C. DeWitt, Editor. 2015, Springer International Publishing: Cham. p. 239–248.

14. DeWitt, J.C., S.J. Blossom, and L.A. Schaider, Exposure to per-fluoroalkyl and polyfluoroalkyl substances leads to immunotoxicity: epidemiological and toxicological evidence. J Expo Sci Environ Epidemiol, 2019. 29(2): p. 148–156.

15. Granum, B., et al., Pre-natal exposure to perfluoroalkyl substances may be associated with altered vaccine antibody levels and immune-related health outcomes in early childhood. 10.3109/1547691X.2012.755580, 2013. 10(4): p. 373–379.

16. Translation, O.o.H.A.a., NTP Monograph on immunotoxicity associated with exposure to perfluorooctanoic acid (PFOA) or perfluorooctane sulfonate (PFOS), U.S.D.o.H.a.H. Services, Editor. 2016.

17. Organization, W.H. WHO Coronavirus (COVID-19) Dashboard. 2023; Available from: https://covid19.who.int.

18. Y, S., et al., COVID-19 infection: the perspectives on immune responses. Cell death and differentiation, 2020. 27(5).

19. Baj, J., et al., COVID-19: Specific and Non-Specific Clinical Manifestations and Symptoms: The Current State of Knowledge. Journal of Clinical Medicine, 2020. 9(6): p. 1753.

20. Tay, M.Z., et al., The trinity of COVID-19: immunity, inflammation and intervention. Nature Reviews Immunology, 2020. 20(6): p. 363–374.

21. Chen, G., et al., Clinical and immunological features of severe and moderate coronavirus disease 2019. The Journal of clinical investigation, 2020. 130(5): p. 2620–2629.

22. García, L.F., Immune response, inflammation, and the clinical spectrum of COVID-19. Frontiers in immunology, 2020. 11: p. 1441.

23. Sly, P.D., et al., The interplay between environmental exposures and COVID-19 risks in the health of children. Environmental Health, 2021. 20(1): p. 1–10.

24. Clerbaux, L.-A., et al., Factors Modulating COVID-19: A Mechanistic Understanding Based on the Adverse Outcome Pathway Framework. Journal of Clinical Medicine, 2022. 11(15): p. 4464.

25. Ahmad, S., Y. Wen, and J.M.K. Irudayaraj, PFOA induces alteration in DNA methylation regulators and SARS-CoV-2 targets Ace2 and Tmprss2 in mouse lung tissues. Toxicology Reports, 2021. 8: p. 1892–1898.

26. Nielsen, C. and A. Jöud, Susceptibility to COVID-19 after High Exposure to Perfluoroalkyl Substances from Contaminated Drinking Water: An Ecological Study from Ronneby, Sweden. International Journal of Environmental Research and Public Health, 2021. 18(20): p. 10702.

27. Ji, J., et al., Association between urinary per- and poly-fluoroalkyl substances and COVID-19 susceptibility. Environment International, 2021. 153: p. 106524.

28. Baumert, B.O., et al., Environmental pollutant risk factors for worse COVID-19 related clinical outcomes in predominately hispanic and latino populations. Environmental Research, 2024. 252: p. 119072.

29. Grandjean, P., et al., Severity of COVID-19 at elevated exposure to perfluorinated alkylates. PLoS One, 2020. 15(12): p. e0244815.

30. Catelan, D., et al., Exposure to perfluoroalkyl substances and mortality for COVID-19: a spatial ecological analysis in the Veneto region (Italy). International journal of environmental research and public health, 2021. 18(5): p. 2734.

31. Enkirch, T. and V. Von Messling, Ferret models of viral pathogenesis. Virology, 2015. 479: p. 259–270.

32. Maher, J.A. and J. DeStefano, The Ferret: An Animal Model to Study Influenza Virus. Lab Animal, 2004. 33(9): p. 50–53.

33. Shou, S., et al., Animal Models for COVID-19: Hamsters, Mouse, Ferret, Mink, Tree Shrew, and Non-human Primates. Frontiers in Microbiology, 2021. 12.

34. Zhao, Y., et al., Ferrets: A powerful model of SARS-CoV-2. Zool Res, 2023. 44(2): p. 323–330.

35. Peng, B., H. Li, and X.-X. Peng, Functional metabolomics: from biomarker discovery to metabolome reprogramming. Protein & Cell, 2015. 6(9): p. 628–637.

36. Edison, A.S., et al., NMR: Unique Strengths That Enhance Modern Metabolomics Research. Analytical Chemistry, 2021. 93(1): p. 478–499.

37. Emwas, A.H., et al., NMR Spectroscopy for Metabolomics Research. Metabolites, 2019. 9(7).

38. Silva, A.V., et al., Associations Between Clinical Signs and Pathological Findings in Toxicity Testing. ALTEX: Alternatives to Animal Experimentation, 2021. 38: p. 198+.

39. Reed, C.E. and S.E. Fenton, Effects of PFOA on Endocrine-Related Systems, in Toxicological Effects of Perfluoroalkyl and Polyfluoroalkyl Substances, J.C. DeWitt, Editor. 2015, Springer International Publishing: Cham. p. 249–264.

40. Bingol, K., et al., Comprehensive Metabolite Identification Strategy Using Multiple Two-Dimensional NMR Spectra of a Complex Mixture Implemented in the COLMARm Web Server. 2016.

41. Cloarec, O., et al., Statistical Total Correlation Spectroscopy: An Exploratory Approach for Latent Biomarker Identification from Metabolic 1H NMR Data Sets. Analytical Chemistry, 2005. 77(5): p. 1282–1289.

42. Garcia-Perez, I., et al., Identifying unknown metabolites using NMR-based metabolic profiling techniques. Nature Protocols, 2020. 15(8): p. 2538–2567.

43. Šidák, Z., Rectangular confidence regions for the means of multivariate normal distributions. Journal of the American statistical association, 1967. 62(318): p. 626–633.

44. Chu, Y.-K., et al., The SARS-CoV ferret model in an infection–challenge study. Virology, 2008. 374(1): p. 151–163.

45. Martina, B.E.E., et al., SARS virus infection of cats and ferrets. Nature, 2025. 425(6961): p. 915-915.

46. Brand, J.M.A.v.d., et al., Pathology of Experimental SARS Coronavirus Infection in Cats and Ferrets. 2008.

47. Kim, Y.-I., et al., Age-dependent pathogenic characteristics of SARS-CoV-2 infection in ferrets. Nature Communications, 2022. 13(1): p. 1–13.

48. JC, D., et al., Perfluorooctanoic acid-induced immunomodulation in adult C57BL/6J or C57BL/6N female mice. Environmental health perspectives, 2008. 116(5).

49. Butenhoff, J.L., et al., Pharmacokinetics of Perfluorooctanoate in Cynomolgus Monkeys. Toxicological Sciences, 2004. 82(2): p. 394–406.

50. Lou, I., et al., Modeling Single and Repeated Dose Pharmacokinetics of PFOA in Mice. Toxicological Sciences, 2008. 107(2): p. 331–341.

51. Hinderliter, P.M., M.P. DeLorme, and G.L. Kennedy, Perfluorooctanoic acid: Relationship between repeated inhalation exposures and plasma PFOA concentration in the rat. Toxicology, 2006. 222(1): p. 80–85.

52. Roberts, M., et al., A Critical Review of Pharmacokinetic Modelling of PFOS and PFOA to Assist in Establishing HGBVs for these Chemicals. Food Standards Australia New Zealand, Majura Park, ACT, Australia, 2016.

53. Jones, P.D., et al., Binding of perfluorinated fatty acids to serum proteins. Environ Toxicol Chem, 2003. 22(11): p. 2639–49.

54. Vanden Heuvel, J.P., et al., Tissue distribution, metabolism, and elimination of perfluorooctanoic acid in male and female rats. J Biochem Toxicol, 1991. 6(2): p. 83–92.

55. Weiss, J.M., et al., Competitive binding of poly- and perfluorinated compounds to the thyroid hormone transport protein transthyretin. Toxicol Sci, 2009. 109(2): p. 206–16.

56. Kannan, K., et al., Perfluorooctanesulfonate and related fluorochemicals in human blood from several countries. Environ Sci Technol, 2004. 38(17): p. 4489–95.

57. Lau, C., Perfluorinated Compounds, in Molecular, Clinical and Environmental Toxicology: Volume 3: Environmental Toxicology, A. Luch, Editor. 2012, Springer Basel: Basel. p. 47–86.

58. Li, K., et al., Molecular mechanisms of PFOA-induced toxicity in animals and humans: Implications for health risks. Environment International, 2017. 99: p. 43–54.

59. DeWitt, J.C., ed. Toxicological Effects of Perfluoroalkyl and Polyfluoroalkyl Substances | SpringerLink. 1 ed. Molecular and Integrative Toxicology. 2015, Humana Cham. 495.

60. Vieira, V.M., et al., Perfluorooctanoic Acid Exposure and Cancer Outcomes in a Contaminated Community: A Geographic Analysis. Environmental Health Perspectives, 2013. 121(3): p. 318–323.

61. Steenland, K. and S. Woskie, Cohort Mortality Study of Workers Exposed to Perfluorooctanoic Acid. American Journal of Epidemiology, 2012. 176(10): p. 909–917.

62. Son, H.-Y., et al., Perfluorooctanoic acid-induced hepatic toxicity following 21-day oral exposure in mice. Archives of toxicology, 2008. 82: p. 239–246.

63. Cui, L., et al., Studies on the Toxicological Effects of PFOA and PFOS on Rats Using Histological Observation and Chemical Analysis. Archives of Environmental Contamination and Toxicology, 2009. 56(2): p. 338–349.

64. Vanden Heuvel, J.P., B.I. Kuslikis, and R.E. Peterson, Covalent binding of perfluorinated fatty acids to proteins in the plasma, liver and testes of rats. Chemico-biological interactions, 1992. 82(3): p. 317–328.

65. Pyper, S.R., et al., PRIC295, a Nuclear Receptor Coactivator, Identified from PPARα-Interacting Cofactor Complex. PPAR research, 2010. 2010(1): p. 173907.

66. Rigamonti, E., G. Chinetti-Gbaguidi, and B. Staels, *Regulation of Macrophage Functions by PPAR-α, PPAR-γ, and LXRs in Mice and Men.* Arteriosclerosis, Thrombosis, and Vascular Biology, 2008. 28(6): p. 1050–1059.

67. Fenton, S.E., et al., Per-and polyfluoroalkyl substance toxicity and human health review: Current state of knowledge and strategies for informing future research. Environmental toxicology and chemistry, 2021. 40(3): p. 606–630.

68. Das, K.P., et al., Perfluoroalkyl acids-induced liver steatosis: Effects on genes controlling lipid homeostasis. Toxicology, 2017. 378: p. 37–52.

69. Di Girolamo, N., 4 - Disorders of the Urinary and Reproductive Systems in Ferrets, in Ferrets, Rabbits, and Rodents (Fourth Edition), K.E. Quesenberry, et al., Editors. 2020, W.B. Saunders: Philadelphia. p. 39–54.

70. Emmett, E.A., et al., Community exposure to perfluorooctanoate: relationships between serum levels and certain health parameters. Journal of occupational and environmental medicine, 2006. 48(8): p. 771–779.

71. Hein, J., et al., Reference ranges for laboratory parameters in ferrets. Veterinary Record, 2012. 171(9): p. 218–218.

72. Carpenter, J.W. and C. Marion, Exotic Animal Formulary-E-Book: Exotic Animal Formulary-E-Book. 2017: Elsevier health sciences.

73. Yang, C.-H., K.P. Glover, and X. Han, Organic anion transporting polypeptide (Oatp) 1a1-mediated perfluorooctanoate transport and evidence for a renal reabsorption mechanism of Oatp1a1 in renal elimination of perfluorocarboxylates in rats. Toxicology letters, 2009. 190(2): p. 163–171.

74. Yang, C.-H., K.P. Glover, and X. Han, Characterization of cellular uptake of perfluorooctanoate via organic anion-transporting polypeptide 1A2, organic anion transporter 4, and urate transporter 1 for their potential roles in mediating human renal reabsorption of perfluorocarboxylates. Toxicological sciences, 2010. 117(2): p. 294–302.

75. Weaver, Y.M., et al., Roles of rat renal organic anion transporters in transporting perfluorinated carboxylates with different chain lengths. Toxicological sciences, 2010. 113(2): p. 305–314.

76. SJ, F., et al., Perfluorooctanoic acid, perfluorooctanesulfonate, and serum lipids in children and adolescents: results from the C8 Health Project. Archives of pediatrics & adolescent medicine, 2010. 164(9).

77. Steenland, K., T. Fletcher, and D.A. Savitz, Epidemiologic Evidence on the Health Effects of Perfluorooctanoic Acid (PFOA). Environmental Health Perspectives, 2010. 118(8): p. 1100–1108.

78. Fletcher, T., et al., Associations between PFOA, PFOS and changes in the expression of genes involved in cholesterol metabolism in humans. Environment International, 2013. 57-58: p. 2-10.

79. Loveless, S.E., et al., Comparative responses of rats and mice exposed to linear/branched, linear, or branched ammonium perfluorooctanoate (APFO). Toxicology, 2006. 220(2-3): p. 203–217.

80. Clampitt, R. and R. Hart, The tissue activities of some diagnostic enzymes in ten mammalian species. Journal of comparative pathology, 1978. 88(4): p. 607–621.

81. Clark, J.M., F.L. Brancati, and A.M. Diehl, The prevalence and etiology of elevated aminotransferase levels in the United States. Official journal of the American College of Gastroenterology| ACG, 2003. 98(5): p. 960–967.

82. Gallo, V., et al., Serum Perfluorooctanoate (PFOA) and Perfluorooctane Sulfonate (PFOS) Concentrations and Liver Function Biomarkers in a Population with Elevated PFOA Exposure. 2012.

83. Sakr, C.J., et al., Cross-sectional study of lipids and liver enzymes related to a serum biomarker of exposure (ammonium perfluorooctanoate or APFO) as part of a general health survey in a cohort of occupationally exposed workers. J Occup Environ Med, 2007. 49(10): p. 1086–96.

84. Jain, R.B. and A. Ducatman, Selective Associations of Recent Low Concentrations of Perfluoroalkyl Substances With Liver Function Biomarkers: NHANES 2011 to 2014 Data on US Adults Aged ≥20 Years. Journal of Occupational and Environmental Medicine, 2019. 61(4): p. 293–302.

85. V, G., et al., Serum perfluorooctanoate (PFOA) and perfluorooctane sulfonate (PFOS) concentrations and liver function biomarkers in a population with elevated PFOA exposure. Environmental health perspectives, 2012. 120(5).

86. Sakr, C.J., et al., Longitudinal study of serum lipids and liver enzymes in workers with occupational exposure to ammonium perfluorooctanoate. J Occup Environ Med, 2007. 49(8): p. 872–9.

87. Zhang, H., et al., Biological responses to perfluorododecanoic acid exposure in rat kidneys as determined by integrated proteomic and metabonomic studies. PLoS One, 2011. 6(6): p. e20862.

88. Peng, S., et al., An integrated metabonomics and transcriptomics approach to understanding metabolic pathway disturbance induced by perfluorooctanoic acid. Journal of pharmaceutical and biomedical analysis, 2013. 86: p. 56–64.

89. A, H., et al., Mechanistic toxicity study of perfluorooctanoic acid in zebrafish suggests mitochondrial dysfunction to play a key role in PFOA toxicity. Chemosphere, 2013. 91(6).

90. Li, X., et al., Lactate metabolism in human health and disease. Signal Transduction and Targeted Therapy, 2022. 7(1): p. 1–22.

91. Dang, E.V., et al., Control of TH17/Treg balance by hypoxia-inducible factor 1. Cell, 2011. 146(5): p. 772–784.

92. Jeong, H.-Y., et al., Anti-inflammatory activity of citric acid-treated wheat germ extract in lipopolysaccharide-stimulated macrophages. Nutrients, 2017. 9(7): p. 730.

93. A, B., et al., Citrate treatment reduces endothelial death and inflammation under hyperglycaemic conditions. Diabetes & vascular disease research, 2012. 9(1).

94. Williams, N.C. and L.A.J. O’Neill, A Role for the Krebs Cycle Intermediate Citrate in Metabolic Reprogramming in Innate Immunity and Inflammation. Front Immunol, 2018. 9: p. 141.

95. Li, T., et al., Longitudinal metabolomics reveals ornithine cycle dysregulation correlates with inflammation and coagulation in COVID-19 severe patients. Frontiers in Microbiology, 2021. 12: p. 723818.

96. Caterino, M., et al., The serum metabolome of moderate and severe COVID-19 patients reflects possible liver alterations involving carbon and nitrogen metabolism. International journal of molecular sciences, 2021. 22(17): p. 9548.

97. Maltais-Payette, I., et al., Association between circulating amino acids and COVID-19 severity. Metabolites, 2023. 13(2): p. 201.

98. Danlos, F.-X., et al., Metabolomic analyses of COVID-19 patients unravel stage-dependent and prognostic biomarkers. Cell Death & Disease, 2021. 12(3): p. 258.

99. Misiura, M. and W. Miltyk, Proline-containing peptides—New insight and implications: A Review. Biofactors, 2019. 45(6): p. 857–866.

100. Ergin Tuncay, M., et al., Modified proline metabolism and Prolidase enzyme in COVID-19. Laboratory Medicine, 2022. 53(5): p. 453–458.

101. Qazi, M.R., et al., High-dose, short-term exposure of mice to perfluorooctanesulfonate (PFOS) or perfluorooctanoate (PFOA) affects the number of circulating neutrophils differently, but enhances the inflammatory responses of macrophages to lipopolysaccharide (LPS) in a similar fashion. Toxicology, 2009. 262(3): p. 207–214.

102. Liang, L., et al., Immunotoxicity mechanisms of perfluorinated compounds PFOA and PFOS. Chemosphere, 2022. 291: p. 132892.

103. Lee, J.W., et al., Integrated analysis of plasma and single immune cells uncovers metabolic changes in individuals with COVID-19. Nature Biotechnology, 2022. 40(1): p. 110–120.

104. Martínez-Gómez, L.E., et al., Metabolic Reprogramming in SARS-CoV-2 Infection Impacts the Outcome of COVID-19 Patients. Frontiers in Immunology, 2022. **Volume** 13 - 2022.

105. Luporini, R.L., et al., Phenylalanine and COVID-19: Tracking disease severity markers. International Immunopharmacology, 2021. 101: p. 108313.

106. Barberis, E., et al., Large-Scale Plasma Analysis Revealed New Mechanisms and Molecules Associated with the Host Response to SARS-CoV-2. Int J Mol Sci, 2020. 21(22).

107. Looker, C., et al., Influenza vaccine response in adults exposed to perfluorooctanoate and perfluorooctanesulfonate. Toxicol Sci, 2014. 138(1): p. 76–88.

108. Sud, M., et al., Metabolomics Workbench: An international repository for metabolomics data and metadata, metabolite standards, protocols, tutorials and training, and analysis tools. Nucleic acids research, 2016. 44(D1): p. D463–D470.

109. Nagana Gowda, G.A. and D. Raftery, Quantitating metabolites in protein precipitated serum using NMR spectroscopy. Anal Chem, 2014. 86(11): p. 5433–40.

110. Gouveia, G.J., et al., Long-Term Metabolomics Reference Material. Analytical Chemistry, 2021. 93(26): p. 9193–9199.

111. Nielsen, N.-P.V., J.M. Carstensen, and J. Smedsgaard, Aligning of single and multiple wavelength chromatographic profiles for chemometric data analysis using correlation optimised warping. Journal of Chromatography A, 1998. 805(1): p. 17–35.

112. Delaglio, F., et al., NMRPipe: a multidimensional spectral processing system based on UNIX pipes. J Biomol NMR, 1995. 6(3): p. 277–93.

113. Ulrich, E.L., et al., BioMagResBank. Nucleic Acids Res, 2008. 36(Database issue): p. D402–8.

114. Wishart, D.S., et al., HMDB: the Human Metabolome Database. Nucleic Acids Res, 2007. 35(Database issue): p. D521–6.

115. Madssen, T.S., et al., Repeated measures ASCA+ for analysis of longitudinal intervention studies with multivariate outcome data | PLOS Computational Biology. PLOS Computational Bioligy, 2021. 17(11).

