## Supplemental Figures S1 - S3, Tables S2 and S4 for "PFOA induced metabolic and immune perturbations in a SARS-2 infection model"

**Figure S1**
**A**

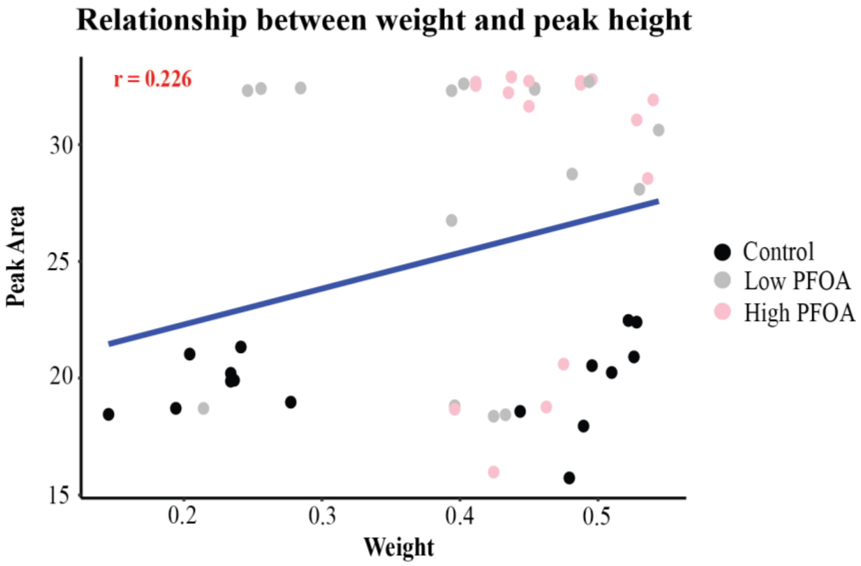


**B**

| | Weight Log2 | | |
|---|---|---|---|
| *Predictors* | *Estimates* | *CI* | *p* |
| (Intercept) | 0.29 | 0.15 – 0.42 | **<0.001** |
| Peak Area log2 | 0.00 | -0.00 – 0.01 | 0.062 |
| Observations | 48 | | |
| $R^2$ / $R^2$ adjusted | 0.074 / 0.054 | | |

**Figure S1: Statistical analysis reveals minimal correlation between PFOA exposure and body weight. Related to Figure 1**
(A) Scatter plot depicting the relationship and correlation between PFOA peak area and ferret weight. Blue line = regression line. r = spearman correlation coefficient.
(B) Linear regression statistics show non-significant association between PFOA levels and body weight (p=0.062, $R^2$ = 0.074).

**Figure S2**

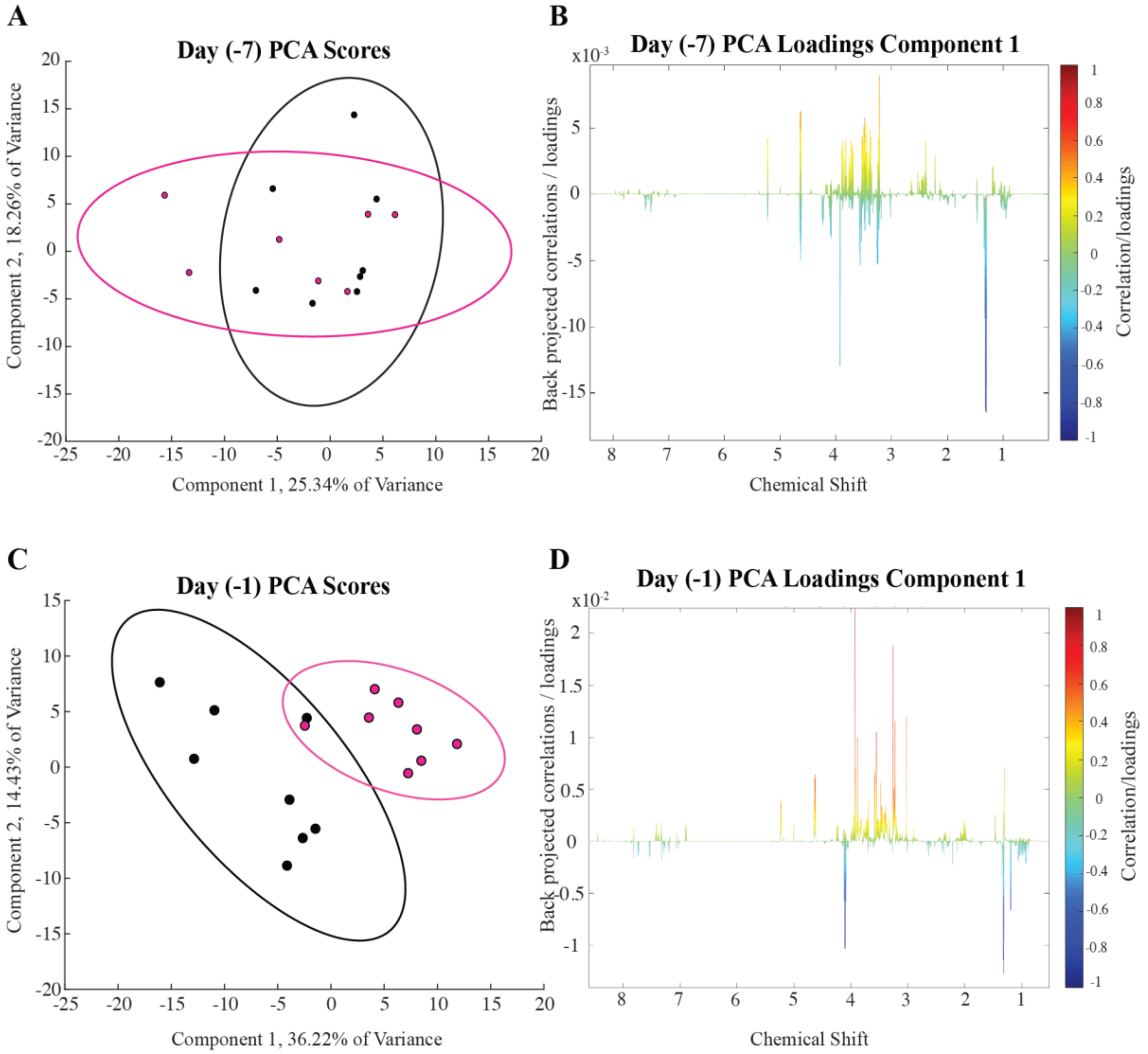


**Figure S2: PCA of -7 DPC and -1 DPC, related to Figure 2**

(A) PCA scores plot of -7 DPC
(B) PCA PC1 Loadings plot of -7 DPC
(C) PCA scores plot of -1 DPC
(D) PCA PC1 loadings plot of -1 DPC

**Figure S3: Renal and Liver Analytes correlation with serum metabolites. Related to figure 2 and Table 1.**

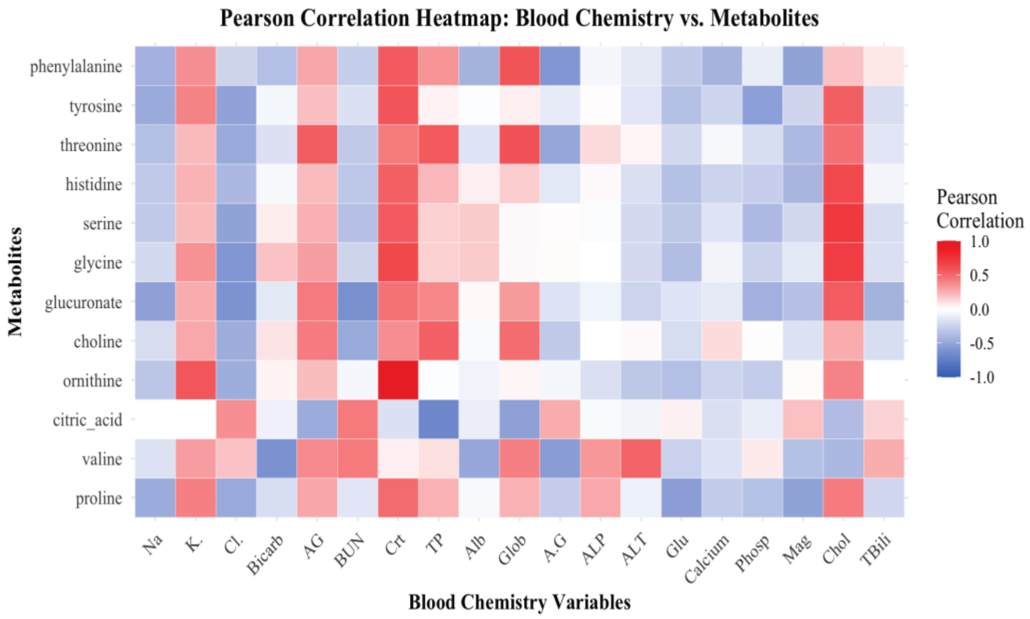


**Figure S3: Renal and Liver Analytes correlation with serum metabolites.**
Hierarchical clustering heatmap of blood chemistry parameters and significantly altered metabolites, revealing functional correlations and group-specific patterns.

**Supplement Tables as separate excel files:**
Table S1 PFOA and Weight analysis, related to Figure 1 – as separate file
Table S3 Blood chemistry analytes analysis, related to Table 1 – as separate file
Table S5 Infection study analysis, related to Figure 4 – as separate file
Table S6 Tissue titer analysis, related to Figure 4 – as separate file

**Table S2** Metabolite Identification table related to Figures 2 and 3

| **Metabolite** | **Database Match** | **KEGG ID** | **PubChem CID** | **Score** |
|---|---|---|---|---|
| **Proline** | bmse000947 | C00148 | 8988 145742 | 3 |
| **Citrate** | bmse000076 | C00158 | 311 | 4 |
| **Ornithine** | bmse000162 | C00077 | 6262 | 3 |
| **Choline** | bmse000285 | C00114 | 305 | 3 |
| **Glucuronate** | bmse000140 | C00191 | 94715 | 4 |
| **Serine** | HMDB03406 | C00740 | 710077 | 3 |
| **Histidine** | bmse001015 | C00135 | 6274 | 3 |
| **Tyrosine** | HMDB00158 | C00082 | 6057 | 3 |
| **Threonine** | HMDB00167 | C00188 | 6288 | 4 |
| **Phenylalanine** | HMDB00159 | C00079 | 6140 | 3 |
| **Isoleucine** | bmse000041 | C00407 | 6306 | 3 |
| **Lactic acid** | bmse000269 | C00256 | 61503 | 4 |
| **Leucine** | bmse000920 | C00123 | 6106 | 3 |
| **Glucose** | HMDB03345 | C00267 | 79025 | 4 |
| **Valine** | HMDB00883 | C00183 | 6287 | 4 |
| **Glycine** | bmse000089 | C00037 | 750 | 4 |

**Table S4**. Generalized Additive Linear Model Results, Related to Figure 3

| Metabolite | Dose_Effect | Time by Control | Time by PFOA |
|---|---|---|---|
| proline | **5.40E-05** | 2.54E-01 | **3.82E-02** |
| citrate | **4.69E-07** | 9.95E-02 | 2.57E-01 |
| ornithine | 3.49E-01 | 3.29E-01 | **8.80E-03** |
| choline | **2.21E-03** | **2.02E-05** | **2.42E-07** |
| glucuronate | **1.22E-08** | 1.84E-01 | **2.09E-02** |
| glycine | **8.62E-11** | 3.53E-01 | 2.58E-01 |
| serine | **9.21E-11** | 2.81E-01 | 4.75E-01 |
| histidine | **1.32E-11** | 2.02E-01 | 4.52E-01 |
| threonine | **2.07E-08** | 7.13E-01 | 9.50E-02 |
| tyrosine | **1.79E-04** | 3.15E-01 | **2.04E-05** |
| phenylalanine | **5.40E-07** | **8.61E-03** | 5.12E-02 |
| isoleucine | 5.91E-01 | **1.26E-02** | 5.14E-01 |
| lactic acid | 3.22E-01 | **6.78E-07** | **4.99E-06** |
| leucine | **1.01E-02** | 2.87E-01 | **1.12E-03** |
| glucose | 2.72E-01 | 5.71E-01 | **1.13E-02** |
